# Cytoplasmic volume and limiting nucleoplasmin scale nuclear size during *Xenopus laevis* development

**DOI:** 10.1101/511451

**Authors:** Pan Chen, Miroslav Tomschik, Katherine Nelson, John Oakey, J. C. Gatlin, Daniel L. Levy

## Abstract

How nuclear size is regulated relative to cell size is a fundamental cell biological question. Reductions in both cell and nuclear sizes during *Xenopus laevis* embryogenesis provide a robust scaling system to study mechanisms of nuclear size regulation. To test if the volume of embryonic cytoplasm is limiting for nuclear growth, we encapsulated gastrula stage embryonic cytoplasm and nuclei in droplets of defined volume using microfluidics. Nuclei grew and reached new steady-state sizes as a function of cytoplasmic volume, supporting a limiting component mechanism of nuclear size control. Through biochemical fractionation, we identified the histone chaperone nucleoplasmin (Npm2) as a putative nuclear size-scaling factor. Cellular amounts of Npm2 decrease over development, and nuclear size was sensitive to Npm2 levels both in vitro and in vivo, affecting nuclear histone levels and chromatin organization. Thus, reductions in cell volume with concomitant decreases in Npm2 amounts represent a developmental mechanism of nuclear size-scaling that may also be relevant to cancers with increased nuclear size.

## INTRODUCTION

Fundamental questions in cell biology concern the regulation of both cell size and sizes of intracellular organelles. The fact that size is regulated within specific ranges suggests that size is important for function (Heald and Cohen-Fix, 2014; Heald and Gibeaux, 2018; Levy and Heald, 2012; Marshall, 2015, 2016). Nuclear size control is of particular interest. There are dramatic reductions in nuclear size during early development (Hara et al., 2013; Jevtic and Levy, 2015), and nuclear size tends to scale with cell size in different species and cell types (Chan and Marshall, 2010; Conklin, 1912; Webster et al., 2009; Wilson, 1925). Furthermore, increased nuclear size is almost uniformly used for cancer diagnosis and prognosis (Chow et al., 2012; Dey, 2010; Jevtic and Levy, 2014; Zink et al., 2004). Nuclear size may therefore play important roles in normal development and cell physiology as well as in disease. Elucidating the functional significance of nuclear size in these various settings requires a mechanistic understanding of the factors and pathways that impinge on the size of the nucleus.

Size control of intracellular structures can be effectively studied in early *Xenopus laevis* embryos. After fertilization, the large 1 mm single cell divides rapidly twelve times without cell growth, giving rise to ~4000 much smaller cells, spanning developmental stages 1-8 (Nieuwkoop and Faber, 1967). Stage 8 coincides with the midblastula transition (MBT), characterized by marked slowing of cell cycles and upregulation of zygotic transcription (Newport and Kirschner, 1982a, b). Gastrulation ensues, encompassing stages 10-12. Between stages 4-8 (i.e. pre-MBT), average cell volume decreases ~160-fold with a concomitant 3.7-fold reduction in nuclear volume; from stage 8-12 (i.e. post-MBT), a more modest 8-fold reduction in cell volume is accompanied by a 3.4-fold reduction in nuclear volume (Jevtic and Levy, 2015). This reproducible scaling of nuclear size provides a robust system with which to characterize and identify mechanisms of nuclear size regulation. Two general models might be invoked to explain how nuclear size scales over development: 1) the expression or localization of developmental regulators of nuclear size may change as development proceeds, and/or 2) the maternal protein pool in the egg contains nuclear assembly or growth components that become limiting as they are partitioned into smaller and smaller cells over development (Goehring and Hyman, 2012; Marshall, 2015; Reber and Goehring, 2015). It is worth noting that these two models are not mutually exclusive and that multiple mechanisms likely contribute to developmental reductions in nuclear size.

To date, primarily developmental regulators of *Xenopus* nuclear size-scaling have been identified. One key mechanism is nucleocytoplasmic transport, mediated by nuclear pore complexes (NPCs) inserted into the nuclear envelope (NE). Classical nuclear import involves the binding of cargos with a nuclear localization signal (NLS) to importin α/β-family karyopherins. Once transported across the NPC, cargos are released by binding of nuclear RanGTP to importin β (Dickmanns et al., 2015; Fried and Kutay, 2003; Madrid and Weis, 2006; Stewart, 2007). Nuclear growth depends on nuclear import (Cox, 1992; Hara et al., 2013), and differences in nuclear import capacity contribute to nuclear size differences in two different *Xenopus* species (Levy and Heald, 2010). Furthermore, during early *X. laevis* embryogenesis, cytoplasmic levels of importin α decrease due to membrane partitioning, leading to reduced nuclear import kinetics and contributing to developmental reductions in nuclear size (Levy and Heald, 2010; Wilbur and Heald, 2013). Importin α cargos important for nuclear growth are nuclear lamins (Jenkins et al., 1993; Newport et al., 1990), NLS-containing intermediate filament proteins that incorporate into the nuclear lamina that underlines the inner nuclear membrane (Dittmer and Misteli, 2011; Wilson and Berk, 2010). Changes in lamin expression levels may also contribute to *Xenopus* developmental nuclear size-scaling (Benavente et al., 1985; Jevtic et al., 2015; Stick and Hausen, 1985). In post-MBT *X. laevis* embryos, redistribution of a population of protein kinase C (PKC) from the cytoplasm to the nucleus leads to phosphorylation-dependent changes in the association of lamins with the NE and concomitant reductions in nuclear size (Edens et al., 2017; Edens and Levy, 2014). Thus, changes in the expression and/or localization of importin α, lamins, and PKC all contribute to developmental nuclear size-scaling in *Xenopus*. Similar factors and mechanisms influence nuclear size in yeast, *C. elegans*, *D. melanogaster*, *T. thermophila*, and cultured human cells (Brandt et al., 2006; Jevtic et al., 2015; Jevtic and Levy, 2014; Kume et al., 2017; Ladouceur et al., 2015; Liu et al., 2000; Malone et al., 2008; Meyerzon et al., 2009; Neumann and Nurse, 2007; Vukovic et al., 2016a; Vukovic et al., 2016b), showing that *Xenopus* development can inform conserved mechanisms of nuclear size control.

Limiting component models have been described for mitotic spindle length scaling (Good et al., 2013; Hazel et al., 2013; Milunovic-Jevtic et al., 2018; Reber et al., 2013) and centrosome size regulation (Decker et al., 2011), however less studied is whether nuclear size might be regulated by limiting components (Goehring and Hyman, 2012; Marshall, 2015; Reber and Goehring, 2015). Microinjection of nuclei into *Xenopus* oocytes resulted in nuclear growth with clustered nuclei growing less (Gurdon, 1976), similar to what has been observed in multinucleate fission yeast cells (Neumann and Nurse, 2007). Consistent with the idea that the amount of surrounding cytoplasm might limit nuclear growth, nuclei assembled in *X. laevis* egg extract grew less when confined in narrow microfluidic channels as opposed to wider channels (Hara and Merten, 2015). This latter study showed that the extent of nuclear growth correlated with the available cytoplasmic space in which interphase microtubule asters could grow, supporting a microtubule-based mechanism for how spatial constraints might limit nuclear growth and steady-state size (Hara and Merten, 2015). Importantly, the aforementioned studies do not fully account for the quantitative reductions in nuclear size observed during early *X. laevis* development, arguing that additional mechanisms must be at work.

Here we test if the volume of embryonic cytoplasm is limiting for nuclear growth, focusing on post-MBT nuclear size-scaling. We encapsulate stage 10 embryonic cytoplasm and endogenous nuclei in droplets in order to precisely manipulate cytoplasmic volume and shape, and we use embryonic cytoplasm as opposed to egg extract to facilitate the identification of bona fide developmental nuclear size regulators. We find that nuclei grow and reach new steady-state sizes as a function of cytoplasmic volume, supporting a limiting component mechanism of nuclear size control. Ruling out candidate limiting components and microtubules, we perform an unbiased biochemical fractionation and identify the histone chaperone nucleoplasmin (Npm2) as a putative nuclear size-scaling factor. Npm2 binds core histones and promotes their assembly into nucleosomes (Frehlick et al., 2007; Platonova et al., 2011). Consistent with Npm2 being a nuclear size-scaling factor, per cell amounts of Npm2 decrease over development and nuclear size is sensitive to Npm2 levels both in vitro and in vivo. We propose that Npm2 influences chromatin topology and/or the composition of the nucleoplasm so as to generate intranuclear forces that promote nuclear growth. In summary, we identify a novel mechanism that contributes to developmental scaling of nuclear size.

## RESULTS

### Cytoplasmic volume contributes to nuclear size-scaling in *Xenopus laevis* embryo extracts

During normal *X. laevis* embryogenesis, individual nuclear volumes scale smaller between the MBT and early gastrulation (stages 8 to 10.5) (Fig. 1A). Because cell sizes also become smaller during this time period, we wondered if cytoplasmic volume might contribute to observed nuclear size-scaling. To test this hypothesis, we isolated extract containing embryonic cytoplasm and endogenous embryonic nuclei from stage 10-10.5 embryos, which have average blastomere volumes of 0.07 nL. We then used microfluidic droplet generating devices to encapsulate this extract in droplets of different volumes and shapes and visualized nuclei by uptake of GFP-NLS (Fig. 1A). After an incubation period, we observed that nuclei grew larger in ~0.8 nL spherical droplets compared to ~0.1 nL spherical droplets (Fig. 1B). In a subset of experiments, we performed live time-lapse imaging of droplets to visualize nuclear growth over time (Movie 1). However in most experiments, we measured nuclear volumes for a large number of droplets at each time point. We then calculated average nuclear and droplet volumes, which we plotted as a function of time. As nuclei grew during incubation, average droplet volumes did not change significantly (Fig. S1A). Nuclei generally reached a new steady-state size after 3-4 hours, consistent with an average cell cycle length of ~4 hours at this stage of development (Duncan and Su, 2004; Murakami et al., 2004). Nuclear volume increased 3.2-fold in ~0.8 nL spherical droplets, but only 1.9-fold in ~0.1 nL spherical droplets (Fig. 1C). In addition, the nuclear growth speed was faster in large droplets compared to small droplets (Fig. S1B). Furthermore, in droplets containing multiple nuclei, the growth of individual nuclei was reduced compared to nuclear growth in similarly sized droplets containing one nucleus (Fig. S1C). These data indicate that the available volume of embryonic cytoplasm can limit nuclear growth.

**Figure 1.**
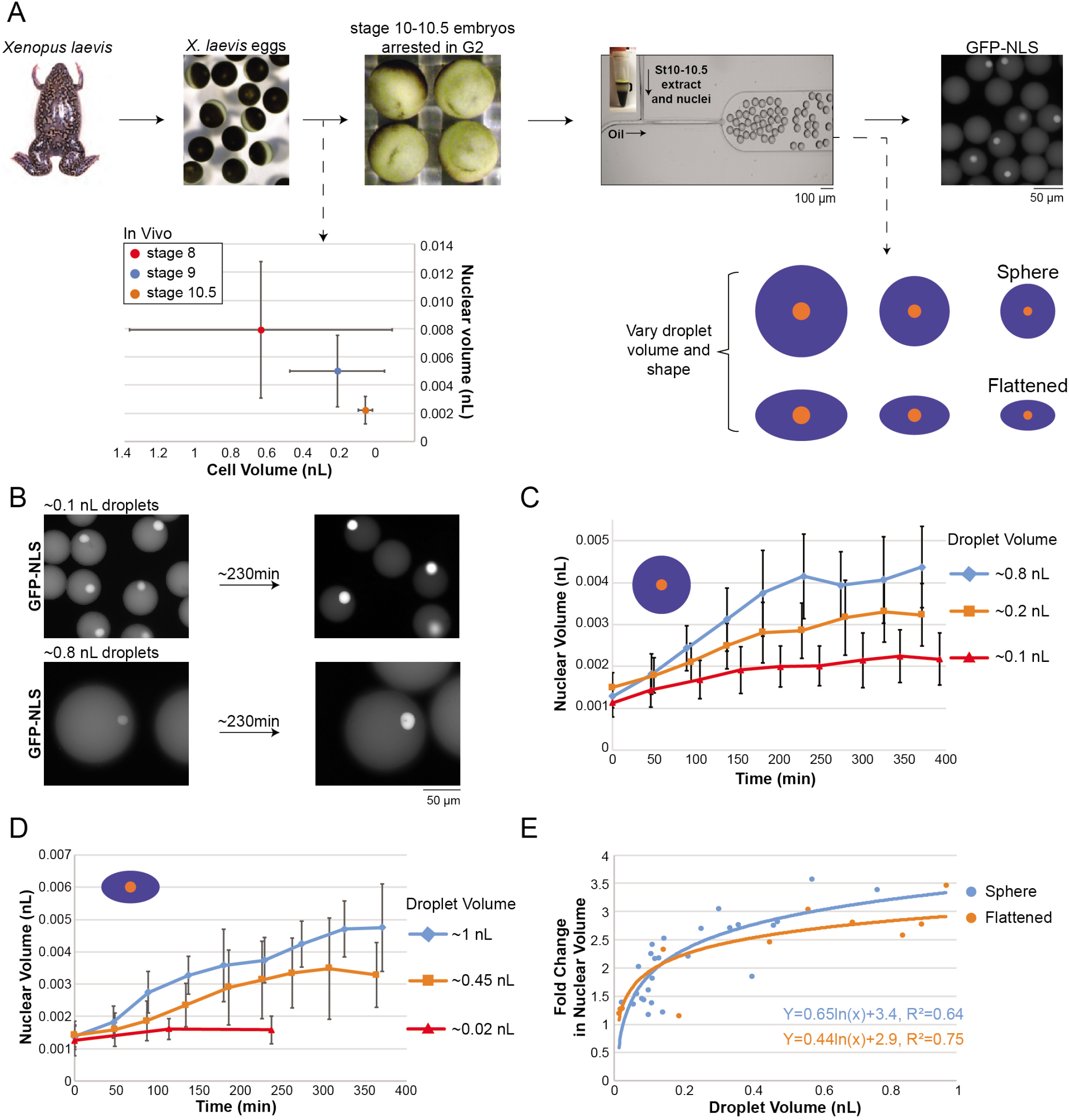
Cytoplasmic volume contributes to nuclear size-scaling in *Xenopus laevis* embryo extracts. **(A)** Top: Schematic diagram of the experimental approach. *Xenopus laevis* eggs were in vitro fertilized. Embryos were allowed to develop to stage 10-10.5 and arrested in late interphase with cycloheximide. Embryonic extract containing endogenous nuclei was encapsulated in droplets using microfluidic devices. Nuclei were visualized by uptake of GFP-NLS. Bottom left panel: Previously generated in vivo nuclear-size scaling data are shown for *X. laevis* stages 8 to 10.5 (Jevtic and Levy, 2015). Bottom right panel: Droplets of differing volumes and shapes can be generated. Blue and orange represent cytoplasm and nuclei, respectively. **(B)** Stage 10 embryo extract and nuclei were encapsulated in different size spherical droplets and incubated at room temperature for 230 minutes. Nuclei were visualized by uptake of GFP-NLS. **(C)** Nuclear volume is plotted as a function of incubation time for spherical droplets of different volumes. At each time point, 18-104 nuclei were quantified (57 nuclei on average). n= 3 extracts. **(D)** Nuclear volume is plotted as a function of incubation time for flattened droplets of different volumes. For flattened droplets, the ratio of the long axis to the short axis was on average ~1.7. At each time point, 10-117 nuclei were quantified (35 nuclei on average). n= 3 extracts. **(E)** Based on the representative nuclear growth curves shown in (C) and (D), for a given droplet size the fold change in nuclear volume was calculated by dividing maximum nuclear volume by initial nuclear volume at t=0. Best-fit logarithmic regression curves are displayed. For each time point of each experiment, 7-311 nuclei were quantified (63 nuclei on average). Data are shown for 26 spherical droplet experiments (blue), 12 flattened droplet experiments (orange), and 23 extracts. All error bars represent standard deviation. See also Figure S1 and Movie 1.

Up to this point, all of our experiments were performed in spherical droplets. Because cell dimensions decrease over development in addition to cell volume, we next tested if nuclear size might be sensitive to cell shape. To test this idea, we generated flattened droplets with a greatly reduced width in one dimension (Fig. 1A). Nuclei still grew in flattened droplets and to a similar extent as in spheres with comparable volumes. In ~1 nL flattened droplets, nuclear volume increased 3.5-fold as opposed to only 1.3-fold in ~0.02 nL flattened droplets (Fig. 1D). To further quantify these effects, we performed a number of experiments using droplet volumes ranging from 0.02 - 1 nL and plotted the fold change in nuclear volume as a function of droplet volume. These data confirm that nuclei reach larger steady-state sizes in larger droplets and that there is no statistically significant difference between spherical and flattened droplets (Figs. 1E and S1D), indicating that in this system cytoplasmic volume, not shape, scales nuclear size.

Previous work demonstrated that microtubule asters sense cytoplasmic volumes in *X. laevis* egg extracts to control nuclear size (Hara and Merten, 2015). To test if microtubules also contribute to nuclear size-scaling in post-MBT embryonic extracts, we encapsulated stage 10 embryo extract treated with nocodazole to depolymerize microtubules. In the absence of microtubules, encapsulated nuclei still grew and to a similar extent as in similarly sized droplets containing microtubules (Fig. S1E). Consistent with these in vitro data, nuclear growth was not reduced when live stage 10 embryos were acutely treated with nocodazole (Fig. S1F). These data indicate that nuclear size-scaling in post-MBT embryos is not microtubule-dependent and suggest that different volume-dependent mechanisms are responsible for nuclear size control in egg extract versus embryos. Taken together, these results demonstrate that nuclear growth and steady-state size scale with cytoplasmic volume in *X. laevis* embryo extracts, independently of cytoplasmic shape and microtubules.

### Cytoplasmic composition contributes to nuclear size-scaling in post-MBT *X. laevis* embryos

We noted that in droplets greater than ~0.5 nL, the nuclear growth speed reached a plateau of 0.012-0.016 pL/min and the change in nuclear volume reached a threshold of 3-3.5-fold (Figs. 1E and S1B). To place this in an in vivo context, we compared nuclear sizes at different developmental stages to our in vitro encapsulation data. When stage 10 extract and nuclei were encapsulated in droplets having similar volumes as cells in stage 8 and 9 embryos, although nuclei grew significantly, they did not reach stage 8 and 9 in vivo nuclear sizes (Fig. 2A). We saw a similar trend when comparing in vitro and in vivo nuclear-to-cytoplasmic (N/C) volume ratios (Fig. S2A). These data suggest that cytoplasmic volume is not sufficient to account for post-MBT nuclear size-scaling. To investigate if changes in cytoplasmic composition might contribute to nuclear size-scaling, we isolated stage 10 nuclei, resuspended them in cytoplasm from earlier embryonic stages, and formed ~0.1 nL droplets. Nuclei in stage 5 cytoplasm grew more than nuclei in stage 6.5 cytoplasm that in turn grew more than nuclei in stage 10 cytoplasm (Figs. 2B-C). Furthermore, when we incubated unencapsulated stage 10 nuclei in bulk egg extract or stage 5 cytoplasm, we also observed significant nuclear growth compared to nuclei incubated in stage 10 cytoplasm (Fig. S2B). These data indicate that differences in the cytoplasmic composition of earlier stage embryos may contribute to nuclear size-scaling in *X. laevis* embryo extracts. One possibility is that these differences result from transcriptional changes that occur at the MBT. Alternatively, because nuclear number increases as development proceeds, components limiting for nuclear growth may become increasingly sequestered into growing numbers of nuclei and concomitantly depleted from the cytoplasm. Thus a limiting component model may explain how both volume and stage-dependent differences in cytoplasmic composition contribute to nuclear-size scaling.

**Figure 2.**
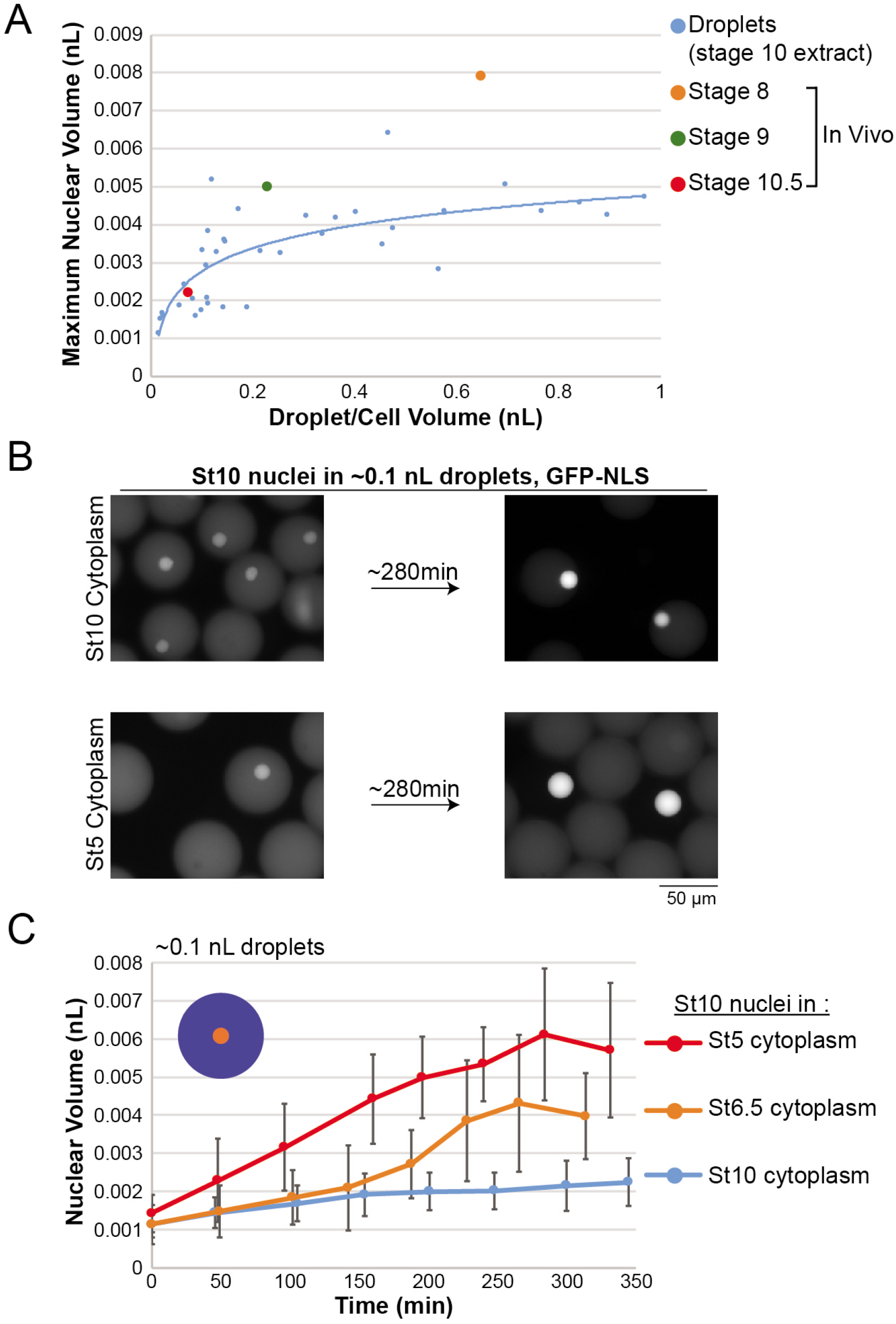
Cytoplasmic composition contributes to nuclear size-scaling in *Xenopus laevis* embryo extracts. **(A)** Based on the data presented in Figure 1C-E, maximum nuclear volume is plotted as a function of droplet volume for stage 10 embryo extract droplets (blue). Also plotted are previously generated in vivo nuclear-size scaling data for *X. laevis* stages 8 to 10.5 (Jevtic and Levy, 2015). **(B)** Stage 10 nuclei were isolated and resuspended in cytoplasmic extract from different embryonic stages. Extracts and nuclei were encapsulated in ~0.1 nL spherical droplets and incubated at room temperature for 280 minutes. Nuclei were visualized by uptake of GFP-NLS. **(C)** Quantification of the experiments described in (B), with nuclear volume plotted as a function of incubation time. At each time point, 10-168 nuclei were quantified (80 nuclei on average). The stage 10 extract data are the same shown in Figure 1C. All error bars represent standard deviation. See also Figure S2.

### Identification of nucleoplasmin as a putative limiting component for nuclear growth

We have shown that the volume of stage 10 cytoplasm is limiting for nuclear growth and that cytoplasm from earlier developmental stages promotes increased nuclear size. These data suggest that one or more cytoplasmic components are limiting for nuclear growth. We reasoned that in droplets of stage 10 extract supplemented with such a limiting component we would observe an increase in nuclear size and upward shift in the nuclear scaling curve. We first took a candidate approach to test if known nuclear size effectors are limiting for growth of stage 10 nuclei. We supplemented stage 10 extract and nuclei with varying concentrations of recombinant lamin B3, lamin B1, or lamin A, formed droplets, and measured the fold change in nuclear size after incubation. Compared to control extract droplets, lamins failed to induce a significant increase in nuclear volume (Fig. S3A). In fact, higher lamin concentrations tended to inhibit nuclear growth as has been previously observed (Jevtic et al., 2015). Using the same approach, we also manipulated PKC activity through a variety of means (Fig. S3B) as well as levels of importin α (data not shown), and in no case did we observe a significant increase in nuclear size. These results suggest that known developmental regulators of nuclear size are not on their own limiting for nuclear growth in restricted cytoplasmic volumes.

Because our candidate approach was unsuccessful, we next undertook a biochemical fractionation approach to identify in an unbiased fashion factors that are limiting for nuclear growth. We reasoned that *X. laevis* egg extract would be a good starting material because: 1) it induces significant growth of stage 10 nuclei (Fig. S2B), 2) it contains large amounts of stockpiled maternal proteins, and 3) it is possible to obtain large volumes of egg extract. We first performed high speed fractionation of egg extract and determined that the nuclear growth inducing activity was present in cytosol but not in the heavy or light membrane fractions (Fig. S3C). We also determined that nuclear growth was dependent on importin α/β mediated import because cytosol treated with the importin β binding domain of importin α (IBB) failed to induce growth of stage 10 nuclei (Fig. S3C). Consistent with this finding, nuclei also failed to grow in stage 10 extract droplets when import was blocked with IBB or wheat germ agglutinin (Fig. S3D). Having determined that the nuclear growth activity was present in egg extract cytosol, we subjected cytosol to fractionation by gel filtration and ion exchange chromatography, monitoring fractions capable of inducing in vitro growth of stage 10 nuclei (Fig. 3A). We ultimately identified two active fractions (Fig. 3B), and we determined the protein composition of these fractions by mass spectrometry (Fig. 3C). Because our nuclear growth activity was dependent on nuclear import (Fig. S3C and S3D), we focused on the most abundant proteins with nuclear localization signals. We were struck by the fact that nucleoplasmin (Npm2) was highly enriched in both fractions (Fig. 3C). Npm2 is a pentameric/decameric histone chaperone and we suspect that it eluted in two different ion exchange chromatography fractions due to its association with different protein complexes. Consistent with this idea, the lower salt elution contained both Npm2 and histones while the higher salt elution contained Npm2 but no core histones (data not show). We verified by western blot using a *Xenopus*-specific antibody the presence of Npm2 in cytosol and fractions active for nuclear growth (Fig. 3D). It is worth noting that several other proteins with histone chaperone activity were identified in the active fractions, three at relatively high abundance (Fig. 3C) and N1/N2, nucleophosmin (Npm1), and Nap1 at lower abundance (data not shown).

**Figure 3.**
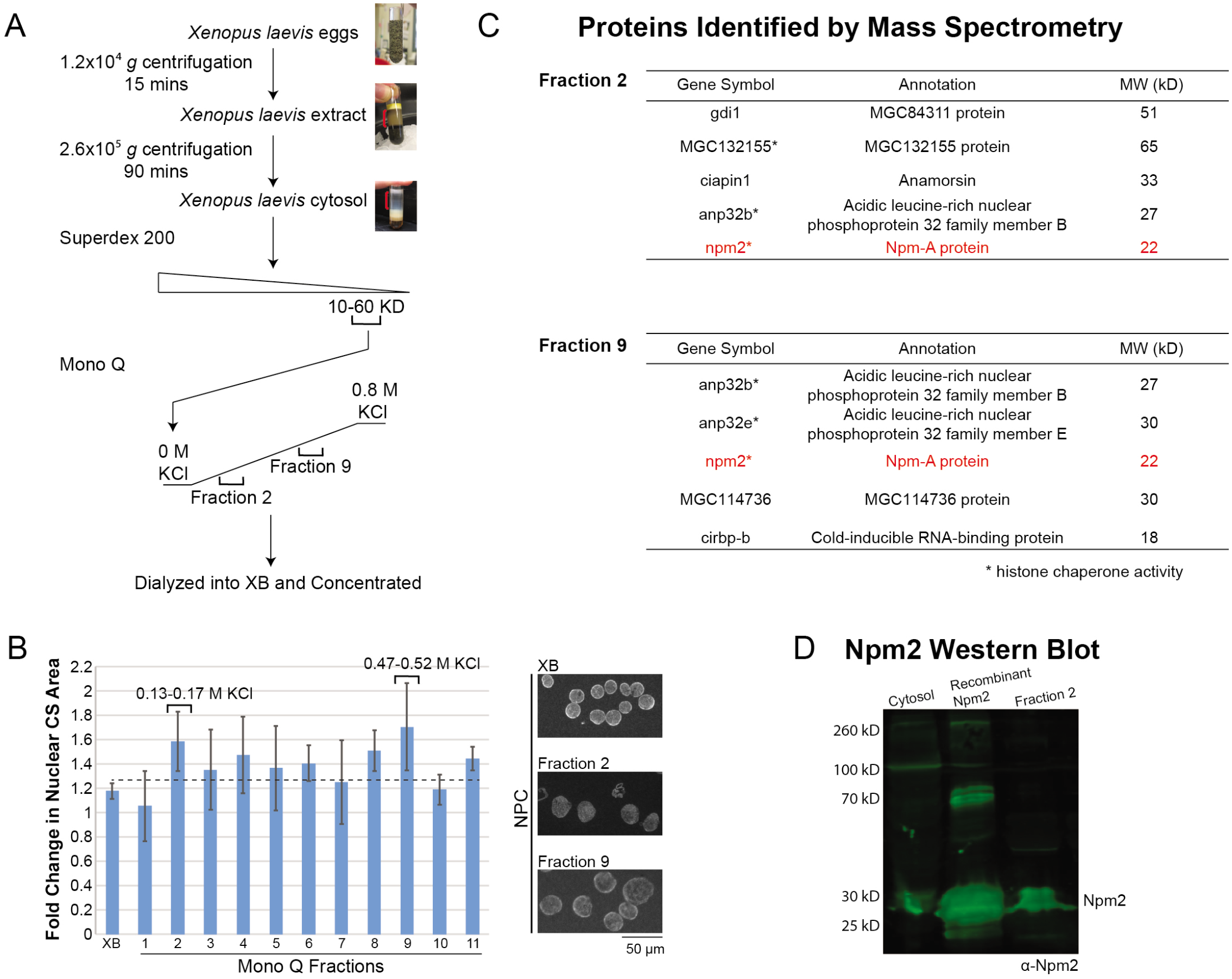
Identification of nucleoplasmin as a putative limiting component for nuclear growth. **(A)** Schematic diagram of the fractionation approach. See Materials and Methods for details. **(B)** Mono Q fractions were dialyzed into XB and concentrated ~10-20-fold. To assay for nuclear growth activity, stage 10 embryo extract and nuclei were supplemented with equivalent volumes of XB or Mono Q fractions. After a 90-minute incubation, nuclei were fixed, spun onto coverslips, and visualized by NPC immunofluorescence with mAb414. Representative images are shown on the right. Nuclear cross-sectional (CS) areas were measured and the fold change was calculated relative to the pre-incubation nuclear size. Means and standard deviations from 3 independent fractionation experiments are shown. At least 160 nuclei were quantified for each condition. The dotted line indicates the top of the error bar for the XB control. We selected fractions with the largest fold changes in nuclear size and with standard deviation error bars above the dotted line, namely fractions 2 and 9. **(C)** Proteins in fractions 2 and 9 were identified by mass spectrometry. The most abundant NLS-containing proteins are shown. Asterisks indicate proteins with histone chaperone activity. **(D)** An anti-Npm2 western blot was performed with 2 μL egg extract cytosol, 5.6 μg purified recombinant Npm2, and 5 μL of fraction 2. All error bars represent standard deviation. See also Figure S3.

### Nucleoplasmin contributes to nuclear size-scaling in vitro

Having identified Npm2 as a putative limiting component for nuclear growth, we next tested if increasing the Npm2 concentration in vitro would increase nuclear size. We first tried supplementing stage 10 extract with varying concentrations of recombinant Npm2 but observed minimal effects on nuclear size (data not shown). We reasoned that Npm2 alone may not be sufficient to induce nuclear growth, so we supplemented stage 10 extract and nuclei with egg extract cytosol (i.e. mixed extract), formed droplets, and monitored nuclear growth. On its own, cytosol induced a modest increase in nuclear size, however the addition of recombinant Npm2 resulted in a significant increase in nuclear growth and steady-state size. In ~0.1 nL droplets, nuclear volume increased to ~3.5 pL in mixed extract supplemented with Npm2, but only reached ~2 pL in mixed extract alone (Fig. 4A). To identify the Npm2 concentration with the maximal effect on nuclear size, we supplemented bulk unencapsulated mixed extract with varying amounts of recombinant Npm2 and determined that 3.5 μM caused the largest increase in nuclear size. We also showed that Npm2 lacking its NLS (i.e. Npm-core) failed to increase nuclear size at 3.5 μM, demonstrating that nuclear import of Npm2 is required to have an effect on nuclear size (Fig. S4A). It is worth noting that the nuclear size increases observed in droplets are much greater than in bulk extract, likely because bulk extract contains a high concentration of nuclei that frequently cluster, something that has previously been shown to constrain nuclear growth (Gurdon, 1976; Hara and Merten, 2015; Neumann and Nurse, 2007).

**Figure 4.**
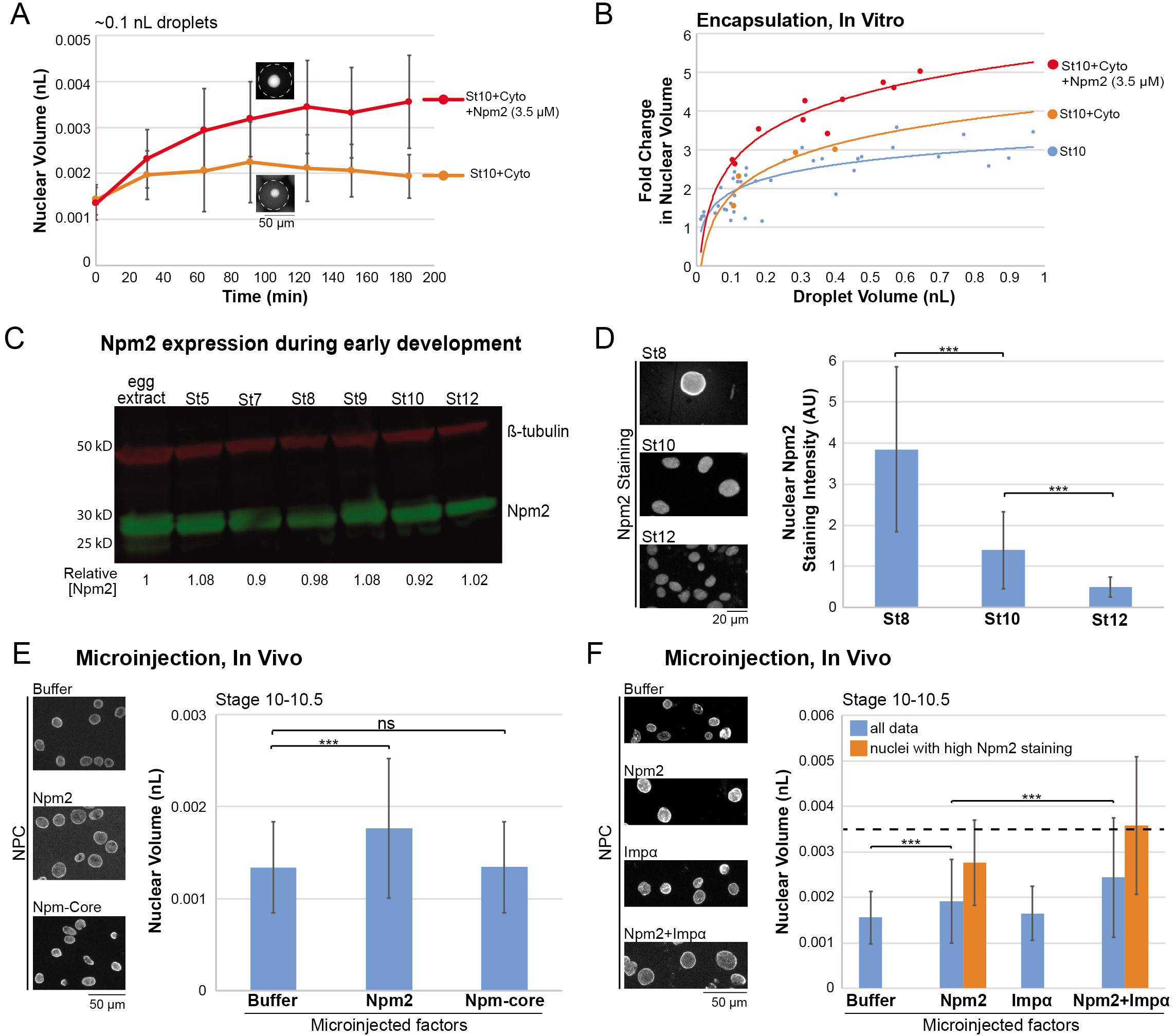
Nucleoplasmin contributes to nuclear size-scaling in vitro and in vivo. **(A)** Stage 10 embryo extract and nuclei were mixed with egg extract cytosol at a 1:4 ratio (St10+Cyto, also called “mixed extract”), a ratio we optimized for nuclear growth (data not shown). Mixed extract was supplemented with 3.5 μM recombinant Npm2 or an equivalent volume of XB. Spherical ~0.1 nL extract droplets were generated and incubated at room temperature as in Figure 1. Nuclei were visualized by uptake of GFPNLS. Nuclear volume is plotted as a function of incubation time. At each time point, 8-182 nuclei were quantified (63 nuclei on average). Representative images are shown at ~90 min. **(B)** Based on the representative nuclear growth curves shown in (A), for a given droplet size the fold change in nuclear volume was calculated by dividing maximum nuclear volume by initial nuclear volume at t=0. Best-fit logarithmic regression curves are displayed. For each time point of each experiment, 8-182 nuclei were quantified (51 nuclei on average). Data from three independent experiments are shown. Droplet data for encapsulated stage 10 extract and nuclei (St10) are the same shown in Figure 1E. **(C)** Extracts were prepared from eggs and different stage embryos. 2 μl of each extract were analyzed by a western blot probed for Npm2 and β-tubulin. Npm2 band intensities were quantified and normalized to β-tubulin levels. Npm2 concentrations were normalized to the amount present in egg extract, and relative Npm2 concentrations are shown below the image. **(D)** Nuclei from different stage embryo extracts were fixed, spun down onto coverslips, and visualized by immunofluorescence using an Npm2 antibody. Images were acquired with the same exposure time. Total nuclear Npm2 staining intensities were quantified for at least 148 nuclei per stage. **(E)** One-cell *X. laevis* embryos were microinjected with equivalent volumes of XB, recombinant Npm2 protein (to increase the Npm2 concentration by 3.5 μM) or recombinant Npm-core protein (3.5 μM final), allowed to develop to stage 10-10.5, and arrested in G2 with cycloheximide. Nuclei isolated from embryos were fixed, spun onto coverslips, and stained with an NPC antibody (mAb414). Nuclear volumes were measured for at least 74 nuclei per condition per experiment. 25 embryos on average were microinjected per condition, and data from four independent experiments are shown. Representative images are shown. **(F)** One-cell *X. laevis* embryos were microinjected with equivalent volumes of XB, recombinant Npm2 protein (to increase the Npm2 concentration by 3.5 μM), importin-α-E mRNA (350 pg total, Imp α), or recombinant Npm2 protein plus importin-α-E mRNA and allowed to develop to stage 10-10.5 without cycloheximide arrest. Importin-α-E is a phosphomimetic version of human importin α2 with reduced affinity for membranes (Levy and Heald, 2010; Wilbur and Heald, 2013). Nuclei isolated from embryos were fixed, spun onto coverslips, and stained with an NPC antibody (mAb414). Nuclear volumes were measured for at least 330 nuclei per condition per experiment. 25 embryos on average were microinjected per condition, and data from two independent experiments are shown. Representative images are shown. Blue bars represent all nuclear size data, while orange bars represent sizes for nuclei with Npm2 staining intensity values greater than one standard deviation above the XB control (see also Figure S4F). The dotted horizontal line at 3.5 pL corresponds to the predicted stage 10 nuclear volume resulting from a 3.5 μM introduction of Npm2 (see also Figure S4D). ***, p≤0.001; ns, not significant. All error bars represent standard deviation. See also Figure S4.

We next generated droplets of differing volumes containing mixed extract with or without added recombinant Npm2 and quantified the fold change in nuclear size after incubation. Compared to stage 10 extract or mixed extract alone, supplementing mixed extract with 3.5 μM Npm2 caused a dramatic upward shift in the nuclear size-scaling curve for all droplet volumes tested (Fig. 4B). We also noted that in mixed extract, nuclei reached their steady-state size faster than in stage 10 extract alone, suggesting a cytosolic activity promotes more rapid nuclear growth (Fig. S4B). These data provide in vitro support for the idea that Npm2 is one factor that limits growth of stage 10 nuclei.

### Nucleoplasmin contributes to nuclear size-scaling in vivo

If Npm2 is a bona fide nuclear size-scaling factor, then we would expect Npm2 to become limiting and for the amount of Npm2 per cell to decrease over the course of normal development. To test this idea, we measured Npm2 levels in embryos ranging from stage 5-12 and found that total concentrations were largely unchanged (Fig. 4C), consistent with a previous proteomic study (Peshkin et al., 2015). Using recombinant Npm2 of known concentration, we measured the in vivo Npm2 concentration to be ~4.2 μM (Fig. S4C). Using this information and known cell volumes in different stage embryos, we determined that the per cell amount of Npm2 should decrease ~9-fold from stage 8-10 (Fig. S4D). Consistent with these calculations, immunofluorescence showed that nuclear Npm2 staining decreases from stage 8-12 by ~8-fold (Fig. 4D). We also confirmed that Npm2 is present in the cytoplasmic fraction of stage 10 extracts (data not shown), explaining the volume-dependent scaling of nuclear size we observed in vitro (Fig. 1). Taken together, these data support the idea that limiting per cell amounts of Npm2 may contribute to developmental nuclear size-scaling.

If Npm2 is a nuclear size-scaling factor, then in vivo nuclear size should be sensitive to Npm2 amounts. To test this idea, we microinjected one-cell *X. laevis* embryos with recombinant Npm2 protein to increase the Npm2 concentration by 3.5 μM, allowed embryos to develop to stage 10, isolated nuclei, and quantified nuclear size. Npm2 microinjection induced a ~1.3-fold increase in nuclear volume relative to buffer-injected controls, and nuclear size was unaffected in embryos microinjected with Npm-core lacking the NLS (Fig. 4E). When we performed Npm2 immunofluorescence, we observed two populations of nuclei isolated from Npm2-microinjected embryos: one group with control level Npm2 staining and one group with ~4-fold more intense Npm2 staining (Fig S4E-F). These data suggested that microinjected Npm2 protein was not evenly distributed across the one-cell embryo. When we quantified nuclei with more intense Npm2 staining, we observed a ~1.8-fold increase in nuclear volume relative to controls (Fig. 4F). We also plotted nuclear volume as a function of Npm2 nuclear staining intensity and observed a strong positive correlation (Fig. S4F). Taken together, these data show that increasing the in vivo concentration of Npm2 is sufficient to increase nuclear size.

Based on the amount of Npm2 microinjected into embryos, the known in vivo Npm2 concentration, and known cell and nuclear sizes at different developmental stages, we predicted that 3.5 μM microinjected Npm2 would produce a stage 10 maximum nuclear volume of 0.0035 nL if Npm2 was the only factor limiting for nuclear growth (Fig. S4D). Because microinjecting Npm2 alone was not sufficient to induce this predicted increase in nuclear volume (Fig. 4E-F), we wondered if other activities might work additively with Npm2. Given known reductions in cytoplasmic importin α levels and nuclear import capacity over development, we microinjected embryos to increase levels of both Npm2 and importin α. While importin α alone had a minimal effect on nuclear size in stage 10 embryos, Npm2 and importin α together induced a ~1.6-fold increase in nuclear volume, and the increase was even larger at ~2.3-fold when considering nuclei with high Npm2 staining intensity (Fig. 4F). Microinjecting importin α along with Npm2 increased nuclear volume by an additional ~30% relative to Npm2 alone, with some nuclear volumes approaching the predicted value of 0.0035 nL. Further increasing the amount of microinjected Npm2 did not lead to additional increases in nuclear size (data not shown), likely due to high intracellular Npm2 concentrations having a dominant negative effect consistent with Npm2 engaging in non-productive import cycles in which it enters the nucleus without bound histones. While other factors likely also contribute to nuclear size reductions in post-MBT embryos, it is striking that Npm2 and importin α are sufficient to account for a significant extent of in vivo nuclear size-scaling. Taken together, these data support the idea that Npm2 is limiting for nuclear growth and contributes to nuclear size-scaling in vivo during development.

### Nucleoplasmin, nuclear histone levels, and chromatin topology

We next wanted to address how Npm2 might promote nuclear growth. Npm2 is a histone chaperone, primarily for histones H2A/H2B but also for histones H3/H4, that facilitates the assembly of nucleosomes on chromatin (Dilworth et al., 1987; Earnshaw et al., 1980; Frehlick et al., 2007; Kleinschmidt et al., 1985; Kleinschmidt et al., 1990; Platonova et al., 2011). Furthermore, chaperone binding has been shown to facilitate nuclear import of histones (Gurard-Levin et al., 2014; Keck and Pemberton, 2013; Mosammaparast et al., 2005; Onikubo and Shechter, 2016). We first wondered if there is a correlation between nuclear histone levels and nuclear size. *Xenopus* egg extract offers a simplified system to ask this question because nuclei are assembled de novo in vitro and can be grown to differing sizes depending on the reaction incubation time. As nuclei grew we observed an increase in nuclear H2B staining intensity, increasing ~9-fold from when intact nuclei first formed to 60 minutes later when nuclear volume had increased ~5-fold (Fig. 5A). Concomitant with this increase in nuclear H2B staining intensity, we observed that the Hoechst-stained chromatin appeared to occupy proportionately less of the nuclear space and to adopt a more heterogeneous distribution throughout the nucleus. To quantify this change in chromatin organization, we measured the chromatin relative area defined as the area occupied by Hoechst-staining DNA divided by the total nuclear cross-sectional area (Baarlink et al., 2017). As nuclei expanded in egg extract, the chromatin relative area decreased by more than 2-fold (Fig. 5A). By analyzing Hoechst intensity line scans across nuclei we also quantified the shift toward a more heterogeneous chromatin topology in larger nuclei (Fig. 5A and S5A). These results are consistent with nuclear growth and chromatin dispersal observed when HeLa nuclei were microinjected into *Xenopus* oocytes (Gurdon, 1976). Taken together, these data suggest that nuclear growth is accompanied by increased nuclear histone accumulation and changes in chromatin organization.

**Figure 5.**
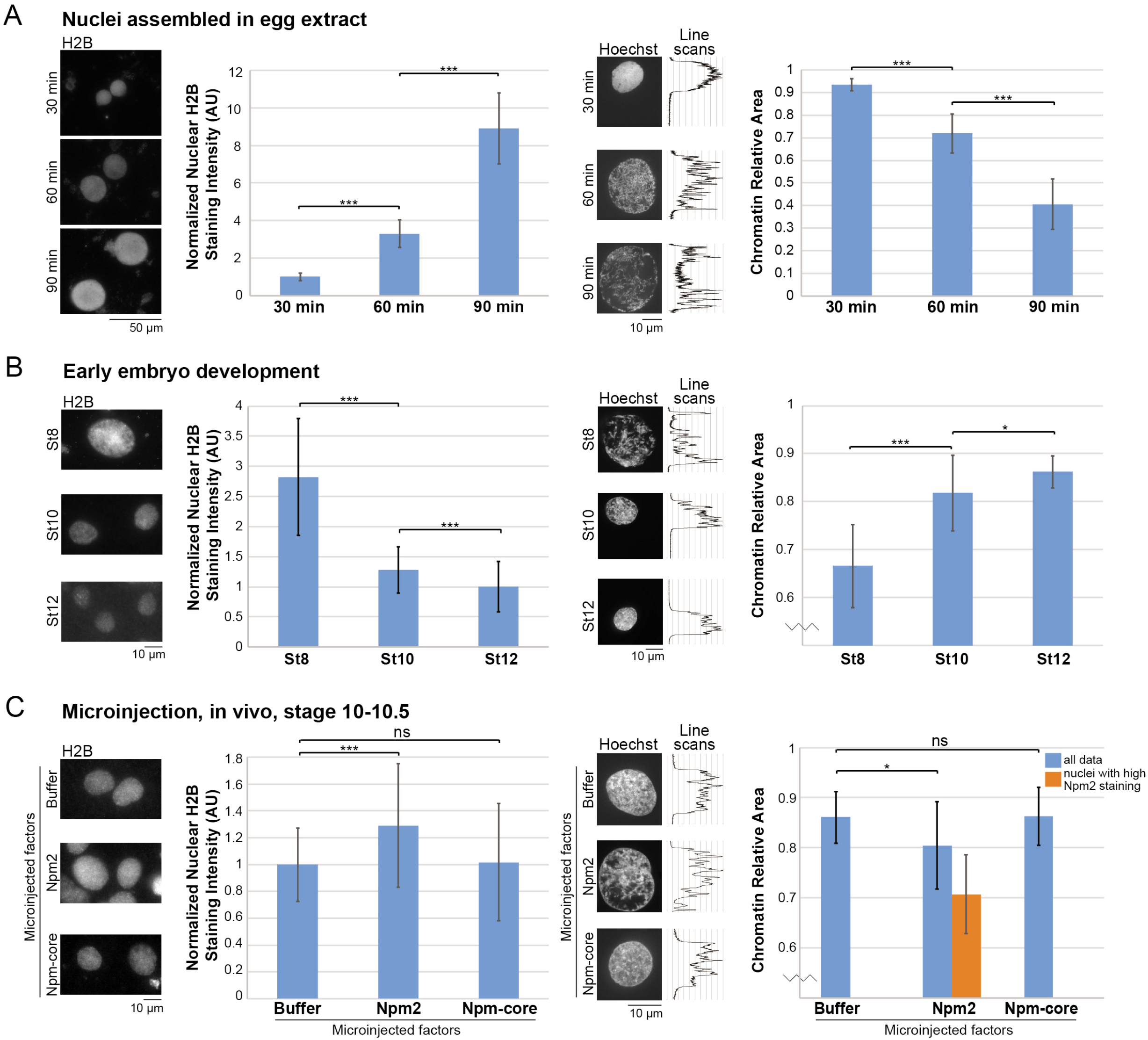
Nucleoplasmin levels correlate with nuclear histone levels and altered chromatin topology. **(A)** Nuclei were assembled de novo in interphasic *X. laevis* egg extract using demembranated *X. laevis* sperm as the chromatin source. After 30, 60, and 90 minutes of incubation, nuclei were fixed, spun down onto coverslips, and visualized by immunofluorescence using an H2B antibody and Hoechst staining. For a given staining approach, images were acquired with the same exposure time. Total nuclear H2B staining intensities were measured for at least 60 nuclei per condition and normalized to the 30-minute time point. To the right of each representative Hoechst image is a Hoechst intensity line scan through the middle of the nucleus. To obtain chromatin relative area measurements, area occupied by Hoechst-staining chromatin was divided by total nuclear area. For chromatin relative area, 23-34 nuclei were quantified per condition (28 nuclei on average). **(B)** Nuclei from different stage embryo extracts were fixed, spun down onto coverslips, and visualized by immunofluorescence using an H2B antibody and Hoechst staining. For a given staining approach, images were acquired with the same exposure time. Total nuclear H2B staining intensities were quantified for at least 100 nuclei per stage and normalized to stage 12. To the right of each representative Hoechst image is a Hoechst intensity line scan through the middle of the nucleus. For chromatin relative area, 14-22 nuclei were quantified per stage (19 nuclei on average). **(C)** One-cell *X. laevis* embryos were microinjected with equivalent volumes of XB, recombinant Npm2 protein (to increase the Npm2 concentration by 3.5 μM), or recombinant Npm-core protein (3.5 μM final), allowed to develop to stage 10-10.5, and arrested in G2 with cycloheximide. Isolated nuclei were fixed, spun down onto coverslips, and visualized by immunofluorescence using an H2B antibody and Hoechst staining. For a given staining approach, images were acquired with the same exposure time. Total nuclear H2B staining intensities were measured for at least 97 nuclei per condition and normalized to the XB-microinjected controls. To the right of each representative Hoechst image is a Hoechst intensity line scan through the middle of the nucleus. For chromatin relative area, 10-63 nuclei were quantified per condition (25 nuclei on average). Data from four independent experiments are shown. Blue bars represent all data, while the orange bar represents data for nuclei with Npm2 staining intensity values greater than one standard deviation above the XB control (see also Figure S4F). ***, p≤0.001; *, p≤0.05; ns, not significant. All error bars represent standard deviation. See also Figure S5.

We next examined whether a similar trend might accompany reductions in nuclear size during development. We already demonstrated that the amount of Npm2 per cell decreases over development, consistent with a potential role as a nuclear size-scaling factor (Figs. 4D and S4D). Along with these reductions in Npm2 amounts, we observed a nearly 3-fold reduction in total nuclear H2B staining intensity from stage 8-12, ~1.3-fold increase in chromatin relative area, and adoption of a more uniform chromatin distribution within the nucleus (Fig. 5B and S5B). These correlations are consistent with the idea that developmental titration of Npm2 leads to reduced nuclear histone levels and an altered chromatin topology that might minimize nuclear growth.

To directly test if Npm2 affects nuclear histone levels and chromatin organization, we revisited our embryo microinjection experiments (Fig. 4E-F). In embryos microinjected to increase Npm2 levels, concomitant with an increase in nuclear size we observed increased nuclear H2B staining intensity, decreased chromatin relative area, and less evenly distributed chromatin throughout the nucleus. None of these effects were observed when Npm-core was microinjected (Fig. 5C and S5C). Consistent with these results, supplementing bulk unencapsulated mixed extract with 3.5 μM recombinant Npm2 also resulted in decreased chromatin relative area, a more heterogeneous chromatin topology (Fig. S5D), and increased nuclear size (Fig. S4A). Because histone chaperones also play important roles in transcriptional regulation (Finn et al., 2012; Frehlick et al., 2007; Gurard-Levin et al., 2014; Onikubo and Shechter, 2016), we tested if the ability of Npm2 to promote nuclear growth was transcription-dependent. Nuclear growth induced by the addition of Npm2 to unencapsulated mixed extract still occurred in the presence of α-amanitin, an inhibitor of RNA polymerase II and III transcription (Seifart and Sekeris, 1969) (Fig. S5E). These data indicate that transcriptional changes are not required for Npm2 to increase nuclear size. Taken together, we propose that Npm2 promotes nuclear growth independently of transcription by increasing nuclear histone levels that in turn impact higher order chromatin structure and distribution within the nucleus.

## DISCUSSION

### Nucleoplasmin is a developmental nuclear size-scaling factor

In this study, by combining microfluidic encapsulation of *X. laevis* embryo extracts and biochemical fractionation, we identified Npm2 as a protein that controls nuclear size. We showed that the per cell amount of Npm2 decreases over *X. laevis* development, as expected for a factor that might become limiting for nuclear growth, and that increasing the in vivo Npm2 concentration is sufficient to increase nuclear size. Nuclei in post-MBT embryos reach a steady-state nuclear size, and data suggest an equilibrium balance of nuclear growth and shrinkage activities that determine this size (Edens et al., 2017; Edens and Levy, 2014; Marshall, 2002). Because per cell Npm2 amounts and nuclear import capacity decrease during post-MBT development, we propose that reductions in these activities reduce nuclear growth and result in the setting of a new equilibrium balance and smaller steady-state nuclear size. Conversely, increasing Npm2 amounts in extract droplets or in vivo enhances nuclear growth, shifting the balance toward larger steady-state nuclear sizes. For these reasons, we propose that Npm2 is a bona fide nuclear size-scaling factor that contributes to developmental reductions in nuclear size. We also believe the identification of Npm2 as a nuclear size effector is novel because it is a nucleoplasmic/volumetric factor, as opposed to previously identified surface area nuclear size regulators that are structural components of the NE, e.g. lamins, lamina-associated proteins, and nesprins (Jevtic and Levy, 2014; Mukherjee et al., 2016; Webster et al., 2009).

Why might Npm2 become physiologically limiting in vivo in post-MBT embryos? In the egg, Npm2 binds to and sequesters maternal core histone stores (Finn et al., 2012; Laskey et al., 1978; Onikubo et al., 2015; Onikubo and Shechter, 2016; Platonova et al., 2011). In pre-MBT embryos, Npm2 and histones are presumably in large excess to the number of nuclei, which explains why supplementing *X. laevis* egg extract with recombinant Npm2 did not increase nuclear size (Levy and Heald, 2010). At the MBT, while the Npm2 concentration remains constant, core histone levels increase (Peshkin et al., 2015; Sun et al., 2014), meaning the Npm2:histone ratio decreases in post-MBT embryos. Furthermore, bulk nuclear import kinetics decrease over development (Levy and Heald, 2010). As a result, limiting amounts of Npm2 are imported more slowly, correlating with reductions in nuclear size and consistent with limiting Npm2 and import both contributing to nuclear size-scaling over development.

Our data support a limiting component model of nuclear size regulation that may be generalizable to nuclear size regulation in other systems. While limiting component models have been used to describe size regulation of other intracellular organelles and structures (Goehring and Hyman, 2012; Marshall, 2015; Reber and Goehring, 2015), we believe this is the first identification of a limiting component that scales nuclear size. Previous work in which *X. laevis* egg extract and de novo assembled nuclei were analyzed in channels of varying dimensions also supported the idea that available cytoplasmic volume scales nuclear growth and steady-state size (Hara and Merten, 2015). Interestingly, in that study nuclear size-scaling was microtubule-dependent, while we find that growth of embryonic nuclei in encapsulated droplets and post-MBT embryos does not dependent on microtubules. This likely represents a difference between eggs and post-MBT embryos and suggests that different mechanisms of nuclear size regulation may dominate depending on developmental stage, as well as perhaps cell type and disease state.

Npm2 does not fully account for nuclear size reductions over development. Our extract encapsulation experiments showed that other cytosolic components were necessary for Npm2 to induce an upward shift in the nuclear size-scaling curve, and increasing Npm2 levels in vivo was not sufficient to achieve nuclear sizes characteristic of pre-MBT developmental stages. It is known that other factors contribute to developmental nuclear size changes, and we did observe additive effects when Npm2 levels were manipulated along with importin α. It is worth noting that increasing importin α levels alone induces a significant increase in nuclear size in pre-MBT embryos (Jevtic and Levy, 2015; Levy and Heald, 2010) but not in post-MBT embryos where both importin α and Npm2 overexpression were required (Fig. 4F). Because Npm2 is a nuclear-imported cargo, these results show that nuclear import kinetics are important throughout early development to scale nuclear size, but that the availability of nuclear sizing cargos may vary over development. In post-MBT embryos, increasing import kinetics alone was not sufficient to increase nuclear size because cargos that drive nuclear growth are limiting, demonstrating that both cargos and transport factors are limiting for nuclear growth in this developmental stage. Thus it is necessary to invoke multiple mechanisms in order to fully account for the reductions in nuclear size that occur during early development. *X. laevis* embryogenesis will no doubt continue to provide insight into these additional mechanisms.

### Mechanism of nucleoplasmin action on nuclear size

By taking an unbiased biochemical fractionation approach, we were able to identify a new factor involved in the regulation of nuclear size that suggests a novel mechanism of nuclear size control. We showed that Npm2 influences nuclear size independently of transcription and that it promotes nuclear localization of histones with concomitant changes in chromatin topology. Open questions remain about precisely how Npm2 promotes nuclear growth. NPC assembly is known to require nucleosome assembly (Inoue and Zhang, 2014; Zierhut et al., 2014), so one possibility is that increased nucleosome assembly by Npm2 leads to larger numbers of NPCs and increased nuclear import capacity. We disfavor this model because growth of stage 10 nuclei with egg cytosol did not lead to an increase in NPC density, and NPC staining intensity was not significantly altered in Npm2-microinjected embryos (data not shown). Another possibility is that increased bulk import of Npm2 and histones leads to passive nuclear enlargement due to increased intranuclear protein content. Nuclei can swell or shrink depending on ionic conditions, pH, temperature, and molecular crowding (Anderson and Wilbur, 1952; Chan et al., 2017; Dahl et al., 2005; Dahl et al., 2004; Enyedi et al., 2016; Hancock and Hadj-Sahraoui, 2009; Korohoda et al., 1968; Kraemer and Coffey, 1970; Pomorski et al., 2000; Stack and Mateyko, 1966). Because histones and Npm2 are highly charged molecules, they could have osmotic effects that promote nuclear expansion. Another model is that Npm2 induces nuclear growth by altering chromatin topology or higher order structure. Npm2 might affect nucleosome assembly, chromatin-association of linker histones and chromatin binding/remodeling proteins, and/or connections between chromatin and the NE (Armiger et al., 2018; Bartholomew, 2014; Chalut et al., 2012; Guelen et al., 2008; Harr et al., 2016; Kruithof et al., 2009; Li and Reinberg, 2011; Luperchio et al., 2014; Macadangdang et al., 2014; Maeshima et al., 2016; Pajerowski et al., 2007; Roopa and Shivashankar, 2006; Schreiner et al., 2015; Shimamoto et al., 2017; Spagnol et al., 2016; Spagnol and Dahl, 2014; Stephens et al., 2017; Stephens et al., 2018; Szerlong and Hansen, 2011; Zidovska et al., 2013). As a result of Npm2-induced alterations in chromatin dynamics and/or higher order organization, chromatin might occupy a larger nuclear volume, potentially providing outward pushing forces that promote nuclear growth.

Common to both the nucleoplasm- and chromatin-mediated models is the idea that outward directed forces from within the nucleus might drive nuclear growth. Such forces could allow for incorporation of lamins and other related proteins into the nuclear lamina that stabilize expansion of the NE, consistent with a recent cell culture study suggesting that nuclear filamentous actin (F-actin) acts in a similar way to promote nuclear growth (Baarlink et al., 2017). Our in vitro system lacks F-actin, so nuclear expansion might rely on other outward pushing forces. Cellular tensegrity (tensional integrity) models have been used to describe cellular architecture. Briefly, balanced tensional and compressive forces mediated by cytoskeletal “struts” and connections to the cell cortex and extracellular matrix help to maintain cell shape (Ingber, 1993, 2003; Ingber et al., 2014; Tadeo et al., 2014). Tensegrity has also been applied to nuclear organization (Aranda-Anzaldo, 2016; Cremer et al., 2000; Maniotis et al., 1997). Invoking such a model, perhaps nuclear size is determined by a balance of forces between chromatin “struts,” nuclear matrix interactions, nucleocytoplasmic osmotic effects, and cytoplasmic connections to the NE via LINC complexes, the cytoskeleton, and the ER (Anderson and Hetzer, 2007, 2008; Cho et al., 2017; Horn, 2014; Lu et al., 2012; Miroshnikova et al., 2017). Changes in the topology or dynamics of chromatin and nuclear F-actin “struts” and/or the biophysical properties of the nucleoplasm might shift the force balance toward smaller or larger steady-state nuclear size. In summary, while extranuclear mechanical forces are known to impact nuclear function (Cho et al., 2017; Kirby and Lammerding, 2018; Miroshnikova et al., 2017; Spagnol et al., 2016), here we propose a role for intranuclear forces in NE expansion.

### Histone chaperones and nuclear size in development and cancer

Our model that Npm2 is a nuclear size-scaling factor over development is consistent with known developmental changes in histone occupancy and transcription. As embryos approach the MBT and total DNA content in the embryo rapidly increases, histone titration is one mechanism that contributes to the upregulation of zygotic transcription (Amodeo et al., 2015; Onikubo and Shechter, 2016). Reduced nucleosome assembly and increased transcription at the MBT correlate with an altered chromatin topology, which by our model is predictive of the reduced nuclear size observed during developmental progression. Mitotic chromosome size scaling is another area of active research (Hara et al., 2016; Hara et al., 2013; Kieserman and Heald, 2011; Ladouceur et al., 2015; Ladouceur et al., 2017; Maresca et al., 2005), and one proposal is that nuclear size and/or nucleocytoplasmic transport of chromosome scaling factors during interphase might determine chromosome size (Heald and Gibeaux, 2018). It is tempting to speculate that limiting nuclear import of Npm2 and histones may represent a coordinated mechanism for the control of nuclear and chromosome size as well as MBT timing. Future work will be required to test these ideas.

While studies in yeast and *Xenopus* have shown that DNA amount is not a primary determinant of nuclear size (Jevtic and Levy, 2014; Jorgensen et al., 2007; Levy and Heald, 2010; Neumann and Nurse, 2007), our data suggest that chromatin organization might influence nuclear growth and steady-state size. Because Npm2 belongs to a large family of nucleophosmin/nucleoplasmin histone chaperones that are conserved throughout metazoans (Finn et al., 2012; Frehlick et al., 2007), it will be important to determine if other histone chaperones impact nuclear size. Interestingly, our biochemical fractionation identified a number of proteins with histone chaperone activity, including N1/N2, nucleophosmin (Npm1), Nap1, and Anp32b/e (Fig. 3C and data not shown). A related area for future investigation will be to examine how nuclear size responds to changes in histone and DNA modifications and recruitment of histone binding proteins that influence the formation of euchromatin and heterochromatin (Audia and Campbell, 2016; Fischle et al., 2003; Grunstein et al., 1995; Harikumar and Meshorer, 2015; Larson et al., 2017; Larson and Narlikar, 2018; Melcer et al., 2012; Nielsen et al., 2001; Stephens et al., 2018; Yu et al., 2011; Zlatanova et al., 2008). It is worth noting that in *T. thermophila* and *Xenopus*, nuclear size is sensitive to the expression of different linker histone H1 isoforms (Freedman and Heald, 2010; Shen et al., 1995).

Whether nuclear size changes in cancer contribute to disease pathology is unknown, and addressing this question necessitates a complete understanding of the mechanisms that control nuclear size. Given that mechanisms of nuclear size regulation identified in *Xenopus* have been shown to act similarly in a variety of other systems, ranging from yeast and *C. elegans* to human cells, we suspect histone chaperones may turn out to represent conserved nuclear size effectors. In support of this idea, 8-cell stage Npm2-null mouse embryos exhibited smaller nuclei compared to wild-type as well as a loss of heterochromatin and altered chromatin organization, with most embryos failing to advance past early embryogenesis (Burns et al., 2003). We find that Npm2 overexpression in vivo in *X. laevis* embryos can induce a ~ 1.3-2.3-fold increase in nuclear volume. Depending on the type of cancer, reported nuclear volume increases range from 1.3-1.7-fold (Abdalla et al., 2009; Mossbacher et al., 1996; Wang et al., 1992), thus the Npm2-induced nuclear size changes we have observed fall within the clinically relevant range of nuclear size changes in cancer. In addition, similar induced changes in nuclear size were sufficient to alter developmental timing programs in *Xenopus* (Jevtic and Levy, 2015, 2017), providing further evidence that nuclear size changes of this magnitude can be physiologically relevant. In addition to previously characterized mechanisms of nuclear size control, our identification of a histone chaperone as a nuclear size effector and proposed chromatin-based model for the regulation of nuclear expansion promise to provide new approaches to address how nuclear size contributes to carcinogenesis.

Histone chaperones have in fact been implicated in cancer. For instance, human Npm1 is frequently overexpressed, mutated, or rearranged in a wide variety of different cancers (Box et al., 2016; Grisendi et al., 2006), as are other histone chaperones (Burgess and Zhang, 2013; Finn et al., 2012; Keck and Pemberton, 2013). This is perhaps not surprising given that mutating nucleosome assembly factors or overexpressing histones can give rise to genome instability and altered gene expression characteristic of many cancers (Groth et al., 2007; Maya Miles et al., 2018; Ransom et al., 2010). It is tempting to speculate that increased histone chaperone expression might contribute to observed nuclear size increases in some cancers, promoting mutagenic potential. Furthermore, histone and DNA modifications are frequently altered in cancer, potentially influencing chromatin organization and nuclear size, and nucleosome structure and histone modifications are possible targets for anti-cancer therapies (Audia and Campbell, 2016; Dawson and Kouzarides, 2012; Gasparian et al., 2011; Lakshmaiah et al., 2014; Safina et al., 2017; Sandoval and Esteller, 2012; Sawan and Herceg, 2010; Waldmann and Schneider, 2013). Future work will address the complex relationships between chromatin structure, nuclear size, and disease pathology, potentially providing new avenues for therapeutic intervention.

## ACKNOWLEDGEMENTS

We thank Kenneth Gerow (University of Wyoming) for help with statistical analysis, Nicolas Blouin and Vikram Chhatre (University of Wyoming) for bioinformatics help identifying NLS-containing proteins, Priscilla Phan for egg extract preparation, David Fay for constructive comments on the manuscript, and members of the Levy, Gatlin, and Oakey labs for helpful advice and discussions. This work was supported by the National Institutes of Health/National Institute of General Medical Sciences (R01GM113028 and P20GM103432), the American Cancer Society (RSG-15-035-01-DDC), and the National Science Foundation Faculty CAREER program (BBBE 1254608).

## AUTHOR CONTRIBUTIONS

Conceptualization, P.C., J.O., J.C.G., D.L.L.; Methodology, K.N. and J.O. designed and fabricated the microfluidic devices; Investigation, P.C. performed all experiments and M.T. performed the fractionations; Writing – Original Draft, P.C., D.L.L.; Writing – Review & Editing, P.C., M.T., K.N., J.O., J.C.G., D.L.L.; Funding Acquisition, J.O., J.C.G., D.L.L.

## MATERIALS AND METHODS

### *Xenopus* embryos, extracts, and microinjections

*X. laevis* embryos were obtained as previously described (Sive et al., 2000). Freshly laid *X. laevis* eggs were in vitro fertilized with crushed *X. laevis* testes. Embryos were dejellied in 3% cysteine (w/v) pH 7.8 dissolved in 1/3x MMR (1x MMR: 0.1 mM EDTA, 0.1 M NaCl, 2 mM KCl, 2 mM CaCl_2_, 1 mM MgCl_2_, 5 mM HEPES pH 7.8). Embryos were developed in 1/3x MMR, staged (Nieuwkoop and Faber, 1967), and arrested in late interphase with 0.15 mg/mL cycloheximide for 60 mins unless otherwise indicated (Lemaitre et al., 1998). Embryo extracts were prepared as previously described (Edens and Levy, 2014, 2016). Briefly, arrested embryos were washed several times in ELB (50 mM KCl, 2.5 mM MgCl_2_, 250 mM sucrose, 10 mM HEPES pH 7.8) containing LPC (10 μg/mL each leupeptin, pepstatin, and chymostatin). For the final wash, the buffer was supplemented with 0.1 mg/mL cytochalasin D and 0.1 mg/mL cycloheximide. The embryos were packed in a tabletop centrifuge at 200 *g* for 1 min at room temperature and excess buffer was removed. The embryos were crushed with a pestle and centrifuged at 10,000 *g* for 10 mins at 16 °C. The cytoplasmic extract containing endogenous embryonic nuclei was collected and supplemented with LPC, 0.02 mg/mL cytochalasin D, 0.1 mg/mL cycloheximide, and energy mix (3.8 mM creatine phosphate disodium, 0.5 mM ATP disodium salt, 0.5 mM MgCl_2_). Extracts were stored on ice until use. In some experiments, stage 10 nuclei were isolated by adding 1 mL ELB to 10-50 μL stage 10 extract, centrifuging at 1600 *g* for 3 minutes to pellet nuclei, and removing the supernatant.

For microinjections, one-cell embryos were dejellied as above thirty minutes after fertilization, transferred to 1/3 MMR containing 2.5% Ficoll (w/v), and microinjected with 10-14 nL volumes using a PicoSprizer III (Parker). To estimate the protein concentration introduced by microinjection, we assumed an egg cytoplasmic volume of 0.5 μL. Microinjected embryos were allowed to develop in 1/3 MMR plus 2.5% Ficoll for 1 hour and then transferred to 1/3 MMR for further development. Extracts from microinjected embryos were prepared as described above, using at least 25 embryos. All *Xenopus* procedures and studies were conducted in compliance with the US Department of Health and Human Services Guide for the Care and Use of Laboratory Animals. Protocols were approved by the University of Wyoming Institutional Animal Care and Use Committee (Assurance #A-3216-01).

### Encapsulation of *Xenopus* extract in microfluidic devices

*X. laevis* embryo extract encapsulation experiments were performed in polydimethylsiloxane (PDMS) microfluidic devices utilizing T-junction droplet generators as previously described (Hazel et al., 2013; Oakey and Gatlin, 2018). PDMS (Sylgard 184, Dow Corning) microfluidic devices were replicated from a negative photoresist-on-silicon master using standard soft lithography protocols (Duffy et al., 1998). Device depth was determined by the thickness to which photoresist was spin-coated upon the silicon wafer. Devices with discrete reservoir depths of 15, 62, 85, and 120 μm were used in this study to achieve increasingly aspherical droplets. PDMS replicas were trimmed and holes were punched using sharpened blunt syringe tips. Prepared devices were exposed to an oxygen plasma (Harrick Plasma) and placed in conformal contact with a glass cover slip. After baking for 10 min at 70°C, an irreversible bond was formed between the PDMS and glass, allowing sealed devices to be used as fluidic networks.

For most experiments, extracts containing cytoplasm and endogenous nuclei were prepared from stage 10-10.5 embryos. To visualize nuclei, extracts were supplemented with 0.04-0.14 mg/mL recombinant GST-GFP-NLS or GST-mCherry-NLS. To limit liquid permeability through the PDMS walls, devices were submerged in ELB for 2 hours prior to encapsulation and during imaging. *X. laevis* embryo extract and carrier oil (Pico-Surf^TM^ 2, 2% in Novec 7500, Dolomite Microfluidics, Cat No 3200282) were loaded into separate syringes and connected to their respective channel inlets via Tygon^®^ microbore tubing (0.010” ID x 0.030” OD, Saint-Gobain Performance Plastics, Cat No AAD04091). Fluid flow to the device was established using a syringe pump (neMYSYS, Cetoni, Kent Scientific, Chemyx). Generally, oil and extract flow rates were 1-10 μl/min and 0.1-1 μl/min, respectively. Relative flow rates were adjusted to vary droplet volume. Filled devices were sealed with acrylic nail polish and droplets were stored in microfluidic reservoirs for imaging. Devices were filled and stored at 4°C prior to imaging at room temperature.

### *Xenopus* egg extract, nuclear assembly, and fractionation

*X. laevis* metaphase-arrested egg extract (Good and Heald, 2018; Maresca and Heald, 2006) and demembranated sperm chromatin (Hazel and Gatlin, 2018; Murray, 1991) were prepared as previously described. Freshly prepared egg extract was supplemented with LPC, cytochalasin D, and energy mix. De novo nuclear assembly was performed as previously described (Chen and Levy, 2018). Interphase-arrested egg extract was generated by supplementation with 0.6 mM CaCl_2_ and 0.15 mg/mL cycloheximide followed by a 25-minute room temperature incubation. A small aliquot of extract was used to test for de novo nuclear assembly, and only those extracts capable of robust nuclear assembly were used for biochemical fractionation. Interphase-arrested egg extracts were centrifuged in a Beckman TL-100 centrifuge for 90 mins at 55,000 rpm using a Beckman TLS-55 rotor at 4°C, to generate cytosol, light membrane, and heavy membrane layers. The cytosol layer was collected and filtered (0.22 μm pore size, 13 mm cellulose acetate syringe filter). 0.8-1.0 mL of filtered cytosol was loaded onto a Superdex 200 10/300 GL column (GE Healthcare) equilibrated in XB (100 mM KCl, 0.1 mM CaCl_2_, 1 mM MgCl_2_, 50 mM sucrose, 10 mM HEPES, pH 7.8). An Akta Purifier FPLC (GE Healthcare) was used to fractionate the cytosol into 40-0.4 mL fractions. Activity assays were performed by supplementing 5 μl of stage 10 embryo extract with 1 μl of each fraction or XB as a control and incubating at room temperature for 90 minutes, followed by nuclear fixation, imaging, and CS area quantification as described under “Immunofluorescence and microscopy.” Active fractions were selected that induced a greater than 14% increase in nuclear CS area relative to XB controls. Active fractions were collected and loaded on a Mono Q 5/50 GL (GE healthcare) anion exchange chromatography column. Bound proteins were eluted using a 0-0.8 M KCl linear gradient in XB over a 6 mL volume, and 0.3 mL fractions were collected. Each fraction was dialyzed against XB buffer for ~2 hours at 4°C (3.5 kD cutoff; Thermo Fisher Scientific, Cat No 69550) and concentrated 10-20-fold using centrifugal filter devices (Microcon YM-10, Millipore, Cat No 42407) at 14,000 *g* for 30-60 mins at 4°C. Activity assays were again performed and active fractions were subjected to mass spectrometry for protein identification. Overall, the fractionation was repeated three separate times with roughly similar activity elution profiles.

### Protein sequence analysis and identification by LC-MS/MS

Active fractions were separated on a 10% SDS-PAGE gel and stained with Coomasie. Gel bands were excised and submitted to the Taplin Mass Spectrometry Facility (Cell Biology Department, Harvard Medical School) for protein identification. Excised gel bands were cut into approximately 1 mm^3^ pieces. Gel pieces were then subjected to a modified in-gel trypsin digestion procedure (Shevchenko et al., 1996). Gel pieces were washed and dehydrated with acetonitrile for 10 minutes followed by removal of acetonitrile. Pieces were then completely dried in a speed-vac. Rehydration of the gel pieces was with 50 mM ammonium bicarbonate solution containing 12.5 ng/μl modified sequencing-grade trypsin (Promega, Madison, WI) at 4°C. After 45 minutes, the excess trypsin solution was removed and replaced with 50 mM ammonium bicarbonate solution to just cover the gel pieces. Samples were then placed in a 37°C room overnight. Peptides were later extracted by removing the ammonium bicarbonate solution, followed by one wash with a solution containing 50% acetonitrile and 1% formic acid. The extracts were then dried in a speed-vac (~1 hr) and stored at 4°C until analysis. Solution samples were reduced with 1 mM DTT (in 50 mM ammonium bicarbonate) for 30 minutes at 60°C. The samples were then cooled to room temperature and 5 mM iodoacetamide (in 50 mM ammonium bicarbonate) was added followed by a 15-minute incubation in the dark at room temperature. DTT was then added to 5 mM to quench the reaction. Sequence grade trypsin was added at 5 ng/μl and digestion carried out overnight at 37°C. The samples were then desalted using an in-house made desalting column.

On the day of analysis the samples were reconstituted in 5-10 μl of HPLC solvent A (2.5% acetonitrile, 0.1% formic acid). A nano-scale reverse-phase HPLC capillary column was created by packing 2.6 μm C18 spherical silica beads into a fused silica capillary (100 μm inner diameter x ~30 cm length) with a flame-drawn tip (Peng and Gygi, 2001). After equilibrating the column each sample was loaded via a Famos auto sampler (LC Packings, San Francisco, CA) onto the column. A gradient was formed and peptides were eluted with increasing concentrations of solvent B (97.5% acetonitrile, 0.1% formic acid). As peptides eluted they were subjected to electrospray ionization and then they entered into an LTQ Orbitrap Velos Pro ion-trap mass spectrometer (Thermo Fisher Scientific, Waltham, MA). Peptides were detected, isolated, and fragmented to produce a tandem mass spectrum of specific fragment ions for each peptide. Peptide sequences (and hence protein identity) were determined by matching protein databases with the acquired fragmentation pattern by the software program, Sequest (Thermo Fisher Scientific, Waltham, MA) (Eng et al., 1994). All databases include a reversed version of all the sequences and the data were filtered to between a one and two percent peptide false discovery rate.

### Recombinant proteins and mRNA

Recombinant GST-GFP-NLS, lamin A, lamin B1, lamin B3, and PKC βII-ΔNPS were expressed and purified as previously described (Edens et al., 2017; Jevtic et al., 2015; Levy and Heald, 2010). The mCherry sequence was amplified from pEmCherry-C2 (a gift from Anne Schlaitz) by PCR and cloned into pMD49 (a gift from Mary Dasso, NIH) at BamHI and EcoRI, replacing the EGFP sequence to generate a bacterial expression construct for GST-mCherry-NLS (pDL94). GST-mCherry-NLS protein was purified similarly to GST-GFP-NLS (Levy and Heald, 2010). Importin α2-E mRNA was synthesized as previously described (Levy and Heald, 2010). Karsten Weis provided the bacterial expression constructs for Npm2, Npm-core, and His-IBB (pKW312, IBB = amino acids 1-65 of hSRP1α), and these proteins were expressed and purified as previously described (Gorlich et al., 1994; Weis et al., 1996).

### Immunofluorescence and microscopy

Immunofluorescence was performed on nuclei isolated from egg or embryo extracts as previously described (Edens and Levy, 2014, 2016). Briefly, extract containing nuclei was mixed with 20 volumes of fix buffer (ELB, 15% glycerol, 2.6% paraformaldehyde), rotated for 15 mins at room temperature, layered over 5 mL cushion buffer (XB, 200 mM sucrose, 25% glycerol), and spun onto 12 mm circular coverslips at 1000x *g* for 15 mins at 16°C. Nuclei on coverslips were post-fixed in cold methanol for 5 mins and rehydrated in PBS-0.1% NP40. Coverslips were blocked with PBS-3% BSA overnight at 4°C, incubated at room temperature for 1 hr each with primary and secondary antibodies diluted in PBS-3% BSA, and stained with 10 μg/mL Hoechst for 5 mins. After each incubation, 6x washes were performed with PBS-0.1% NP40. Coverslips were mounted in Vectashield mounting medium (Vector Laboratories, Cat No H-1000) onto glass slides and sealed with nail polish. Primary antibodies included mAb414 (BioLegend # 902901, mouse, 1:1000) that recognizes NPC FG-repeats, histone H2B antibody (Millipore # 07-371, rabbit, 1:100) and *X. laevis* Npm2 nucleoplasmin antibody (a gift from David Shechter, Albert Einstein College of Medicine, rabbit, 1:1000). Secondary antibodies included 1:1000 dilutions of Alexa Fluor 488 and 568 anti-mouse IgG (Molecular Probes, A-11001 and A-11004) and Alexa Fluor 488 and 568 anti-rabbit IgG (Molecular Probes, A-11008 and A-11011).

Wide-field microscopy was performed using an Olympus BX63 upright wide-field epifluorescence microscope. This system is equipped to perform multimode, time-lapse imaging using an X-Cite 120LED illumination system. Image acquisition was with a high-resolution Hamamatsu ORCA-Flash4.0 digital CMOS camera at room temperature. Olympus objectives included the PLanApoN 2X (NA 0.08, air), UPLanFLN 20X (NA 0.5, air), and UPLanSApo 40X (NA 1.25, silicon oil). X-Y and Z positions were controlled by a fully motorized Olympus stage. Acquisition and automation were controlled by Olympus cellSens imaging software, and image analysis was performed using Metamorph software (Molecular Devices). Images for measuring fluorescence intensity were acquired using the same exposure times. Total fluorescence intensity and cross sectional nuclear area were measured from original thresholded images using Metamorph software. For nuclear size measurements, live nuclei in droplets were visualized with GFP-NLS or mCherry-NLS while fixed nuclei spun onto coverslips were visualized with mAb414. Nuclear and droplet volumes were extrapolated from cross sectional (CS) area measurements, as previous data showed that CS area accurately predicts total nuclear surface area and volume as measured from confocal z-stacks (Edens and Levy, 2014; Jevtic and Levy, 2015; Levy and Heald, 2010; Vukovic et al., 2016b). For publication, images were cropped using ImageJ but were otherwise unaltered.

Confocal imaging was performed on a spinning-disk confocal microscope based on an Olympus IX81 microscope stand equipped with a five line LMM5 laser launch (Spectral Applied Research) and Yokogawa CSU-X1 spinning-disk head. Confocal images were acquired with an EM-CCD camera (ImagEM, Hamamatsu). Z-axis focus was controlled using a piezo Pi-Foc (Physik Instrumentes), and multiposition imaging was achieved using a motorized Ludl stage. Olympus objective PLanApo 100X (NA 1.4, oil) was used. Image acquisition and all system components were controlled using Metamorph software. To measure chromatin relative area, confocal cross-sectional images of Hoechst-stained nuclei were acquired. Using MetaMorph, thresholding was applied to obtain the area of only the Hoechst-stained chromatin, then less stringent thresholding was applied to obtain total nuclear CS area. Finally, chromatin relative area was calculated by dividing the area occupied by Hoechst-staining chromatin by total nuclear area. To quantify the distribution of chromatin within the nucleus, we acquired two Hoechst intensity line scans through the middle of each nucleus using ImageJ. We reasoned that intensity values along these lines would vary greatly for a heterogeneous chromatin distribution and show less variability for a more uniform chromatin distribution. To measure this variability, the standard deviation of all intensity values along each line was determined and normalized to the average intensity to obtain a value we term the “chromatin heterogeneity index.” Larger values correspond to a more heterogeneous chromatin distribution.

### Western blots

Protein samples (e.g. extract, cytosol, fractionated extract, purified Npm2) were supplemented with SDS-PAGE loading buffer (0.05% Bromophenol blue, 0.1 M DTT, 10% glycerol, 2% SDS, 0.05 M Tris-Cl, pH 6.8) and boiled for 10 mins. Proteins were separated on 12% SDS-PAGE gels and transferred to Immobilon-FL PVDF membrane (Millipore, Cat No IPFL00010) using a tank blotting apparatus (Bio-Rad Laboratories). The membrane was blocked in Odyssey Blocking Buffer (Li-COR Biosciences, Cat No 927-40000) for 60 mins at room temperature, and probed with primary antibodies diluted in Odyssey Blocking Buffer/PBST (1:3) overnight at 4°C. 3-10 minute washes were performed in PBST. Membranes were then probed with secondary antibodies diluted in Odyssey Blocking Buffer supplemented with 0.01% SDS and 0.1% Tween 20 for 60 mins at room temperature. 3-10 minute washes were performed in PBST. Membranes were then rinsed in water and scanned on an Odyssey CLx instrument (Li-COR Biosciences). Band intensities were quantified using Odyssey software (ImageStudio). Primary antibodies included *X. laevis* Npm2 nucleoplasmin antibody (a gift from David Shechter, Albert Einstein College of Medicine, rabbit, 1:1000) and β-tubulin antibody (Santa Cruz Biotechnology, Cat No sc-58884, mouse, 1:100). Secondary antibodies used at 1:20,000 were IRDye 800CW anti-rabbit (Li-Cor 926-32211) and IRDye 680RD anti-mouse (Li-Cor 925-68070).

### Statistical analysis

Averaging and statistical analysis were performed for independently repeated experiments. Where indicated, nuclear size and intensity measurements were normalized to controls. Two-tailed Student’s t-tests assuming equal variances were performed with Minitab 18 or Prism 6 to evaluate statistical significance. For each immunofluorescence coverslip, generally more than 100 nuclei were quantified. The p-values, number of independent experiments, number of nuclei quantified, and error bars are denoted in the figure legends.

---

**Figure S1.**
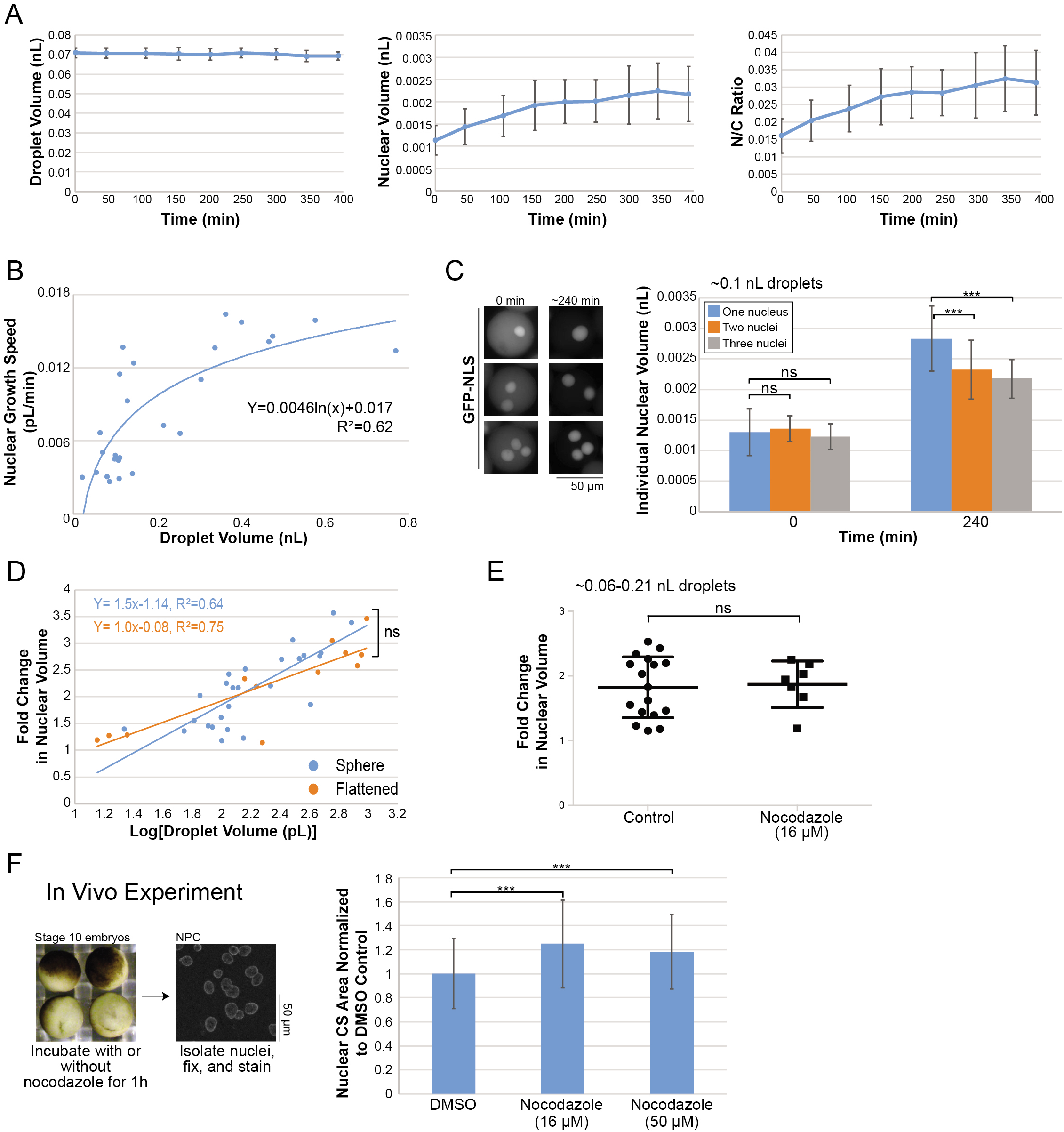
Further characterization of the contribution of cytoplasmic volume to nuclear size-scaling. Related to Figure 1. **(A)** Raw data for one of the experiments shown in Figure 1C. Left panel: droplet volume is plotted as a function of incubation time. Middle panel: nuclear volume is plotted as a function of incubation time. Right panel: Nuclear-to-cytoplasmic (N/C) volume ratio is plotted as a function of incubation time. At each time point, 31-104 nuclei were quantified (53 nuclei on average). **(B)** Based on the representative nuclear growth curves shown in Figure 1C, nuclear growth speeds were calculated as follows: [Nuclear volume (pL) _t1=~150 min_-Nuclear volume (pL) _t=0 min_]/time(t1). A best-fit logarithmic regression curve is displayed. For each experiment and time point, 7-311 nuclei were quantified (72 nuclei on average). Data are shown for 26 spherical droplet experiments and 17 extracts. **(C)** Stage 10 embryo extract and nuclei were encapsulated in ~0.1 nL spherical droplets and incubated at room temperature for 240 minutes. Droplets with differing numbers of nuclei were selected for analysis. Nuclei were visualized by uptake of GFP-NLS, and representative images are shown. Individual nuclear volumes were quantified. For each condition, 57-117 nuclei were quantified (74 nuclei on average). **(D)** The same data presented in Figure 1E are plotted, here as the logarithm of droplet volume in order to facilitate statistical comparison of the sphere and flattened droplet data. Best-fit linear regression lines are displayed. Plotting of nuclear volume versus droplet volume on a log-log plot produced a linear regression line with slope 0.31, indicative of hypoallometric scaling similar to that observed in vivo for nuclear volume versus blastomere volume from stages 3-12 with slope 0.32 (Jevtic and Levy, 2015; Uppaluri et al., 2016). **(E)** Stage 10 embryo extract was supplemented with 16 μM nocodazole. In some control experiments, an equivalent volume of DMSO was added. Nuclear growth in ~0.06-0.21 nL droplets was performed as before. For each time point of each experiment, 7-601 nuclei were quantified (90 nuclei on average). Nocodazole-induced depolymerization of microtubules was verified by imaging fluorescently-labeled tubulin supplemented into extract (data not shown). n=3 extracts. The longer horizontal lines represent the means and the shorter horizontal lines correspond to the standard deviations. Some of the control data are the same as presented in Figure 1E. **(F)** Left panel: The experimental approach is shown. Stage 10-10.5 embryos were incubated with or without nocodazole for 60 mins. For both conditions, cycloheximide was included to arrest embryos in late G2. Nuclei in embryo extracts were fixed, spun down onto coverslips, and visualized by NPC immunofluorescence using mAb414. Right panel: Nuclear CS areas were measured, averaged, and normalized to the mean of the control. For each condition, at least 110 nuclei were quantified. Data from two independent experiments are shown. ***, p≤0.001; ns, not significant. All error bars represent standard deviation.

**Figure S2.**
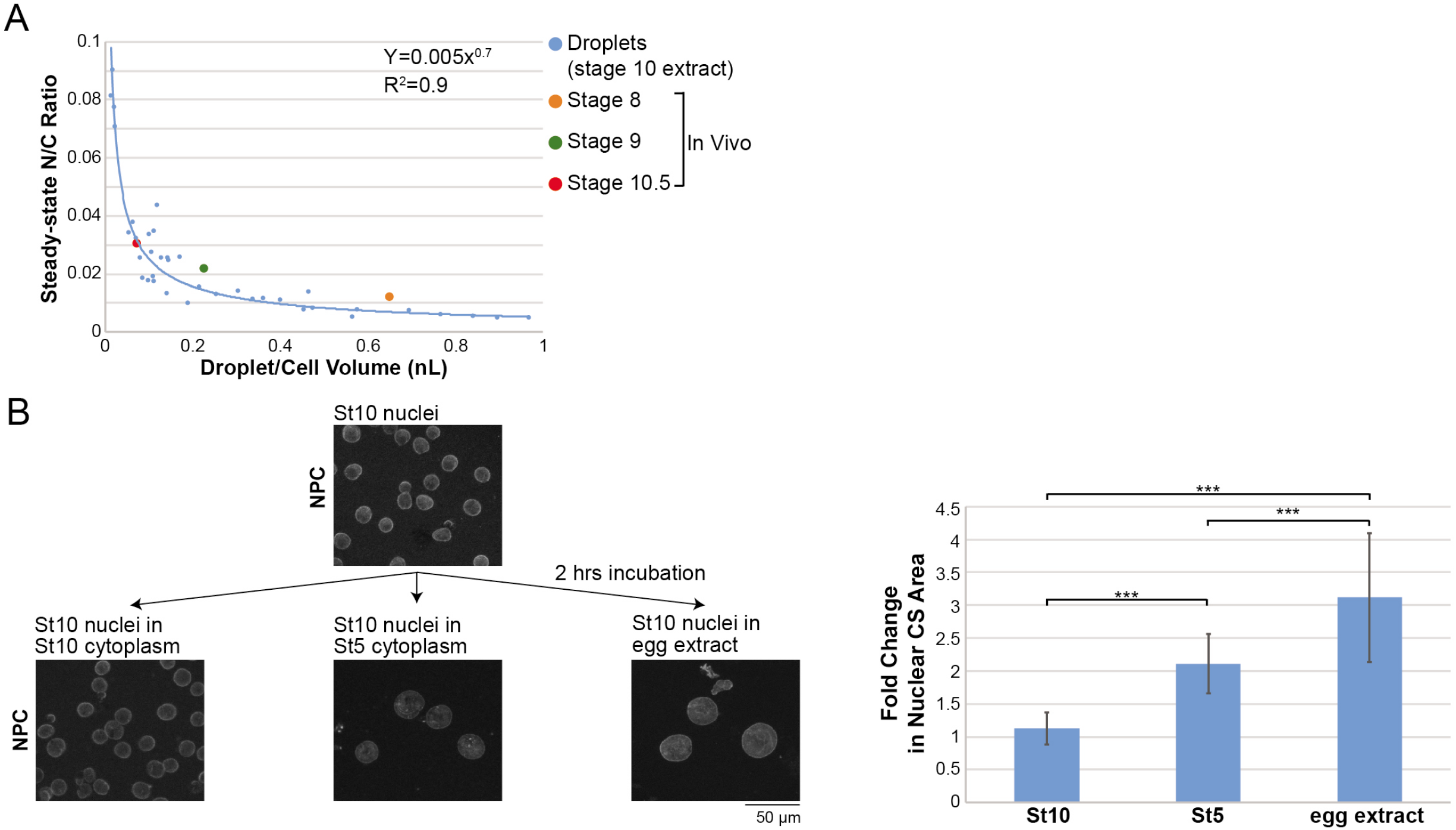
Stage 10 nuclei grow more in extracts from pre-MBT embryos and eggs compared to post-MBT embryo extract. Related to Figure 2. **(A)** Based on the data shown in Figure 1C-E, nuclear-to-cytoplasmic (N/C) volume ratios were calculated by dividing maximum nuclear volume by droplet volume. A best-fit power regression curve is displayed for the droplet data. Also plotted are previously reported in vivo N/C volume ratios for stage 8 to 10.5 embryos (Jevtic and Levy, 2015). **(B)** Stage 10 extract and nuclei were mixed with stage 5 cytoplasmic extract or egg extract at a 1:10 ratio. After a 2-hour incubation, nuclei were fixed, spun down onto coverslips, and visualized by NPC immunofluorescence using mAb414. Nuclear CS areas were measured and the fold change was calculated relative to the pre-incubation nuclear size. At least 700 nuclei were quantified for each condition. Data from two independent experiments are shown. ***, p≤0.001. All error bars represent standard deviation.

**Figure S3.**
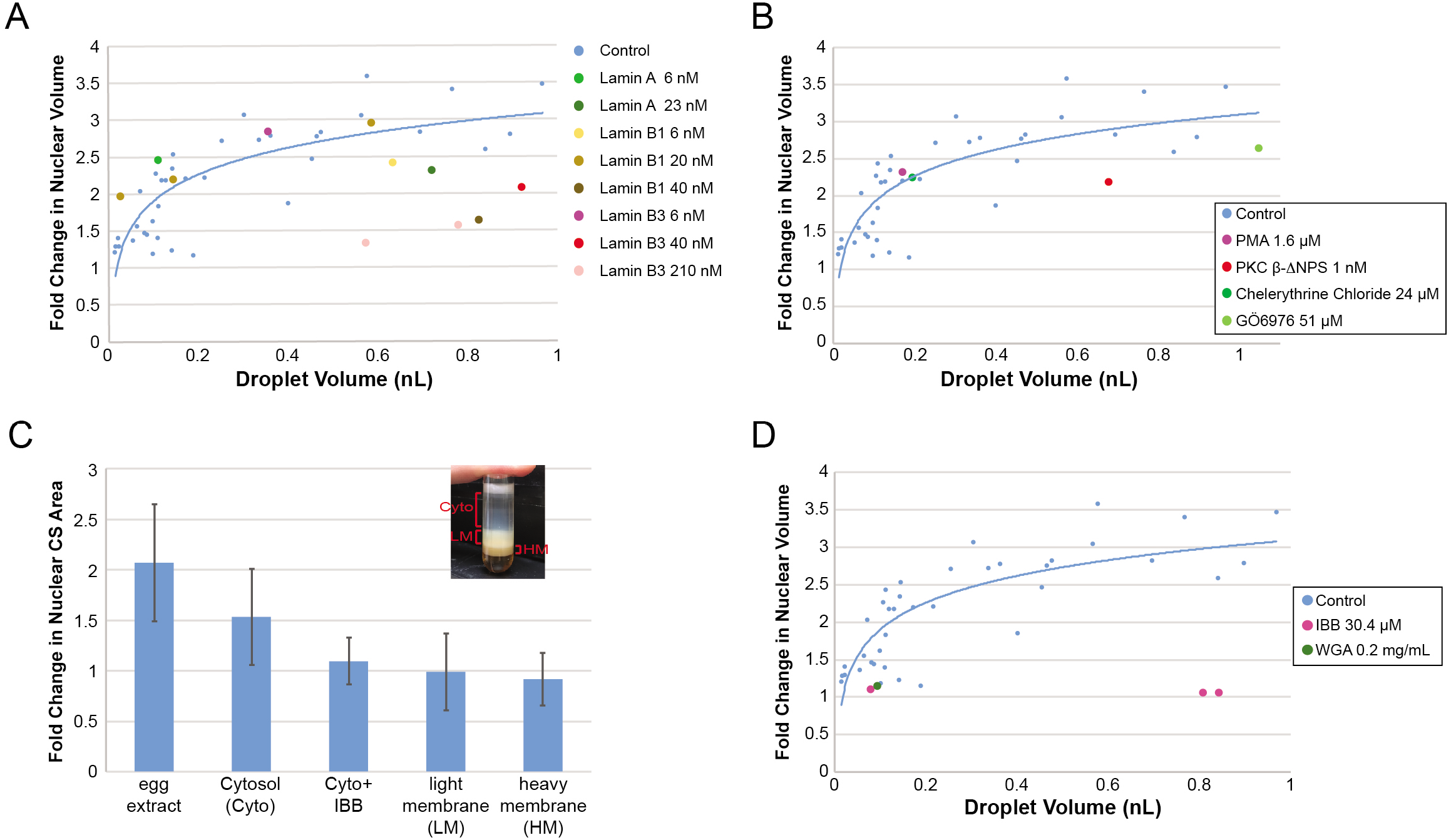
Testing candidate factors and activities that may be limiting for nuclear growth. Related to Figure 3. **(A)** Experiments were performed as in Figure 1C-E except that extracts were supplemented with the indicated concentrations of recombinant nuclear lamin proteins. For each lamin experiment, 8-148 nuclei were quantified at each time point (41 nuclei on average). n=4 extracts. The control data are the same shown in Figure 1E. **(B)** Experiments were performed as in Figure 1C-E except that extracts were supplemented with the indicated reagents. Phorbol 12-myristate 13-acetate (PMA) is a PKC activator, PKC β-ΔNPS is a constitutively active form of PKC β, and chelerythrine chloride and Gö6976 are PKC inhibitors (Edens et al., 2017; Edens and Levy, 2014). For each PKC experiment, 13-49 nuclei were quantified at each time point (32 nuclei on average). n= 2 extracts. The control data are the same shown in Figure 1E. **(C)** *X. laevis* egg extract was subjected to high speed centrifugation to separate the extract into cytosol (Cyto), light membrane (LM), and heavy membrane (HM), as shown in the inset. Nuclei isolated from stage 10 embryo extract were incubated in equivalent volumes of egg extract, cytosol, light membrane, or heavy membrane for 120 mins. For the “Cyto+IBB” experiment, cytosol was supplemented with 30.4 μM IBB prior to incubation. IBB is the importin β binding domain of importin α and inhibits nuclear import (Weis et al., 1996). Nuclei were fixed, spun down onto coverslips, and stained with an antibody against the NPC (mAb414). Nuclear CS areas were measured for at least 200 nuclei per condition and normalized to the pre-incubation nuclear size. Data from two independent experiments are shown. **(D)** Experiments were performed as in Figure 1C-E except that extracts were supplemented with IBB or wheat germ agglutinin (WGA) to block nuclear import (Cox, 1992). For each experiment, 7-212 nuclei were quantified at each time point (53 nuclei on average). n= 3 extracts. The control data are the same shown in Figure 1E. All error bars represent standard deviation.

**Figure S4.**
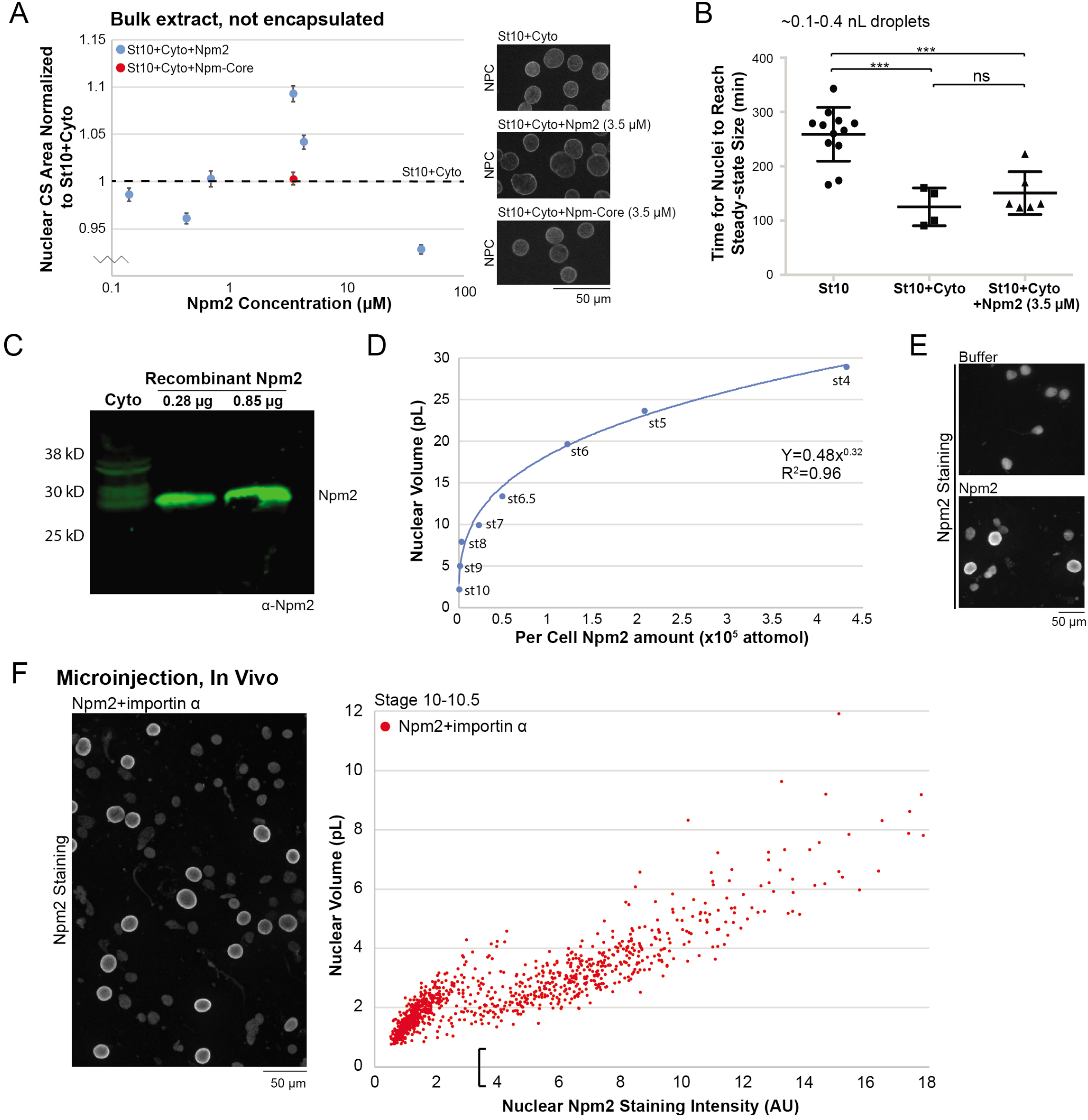
Further characterization of the contribution of nucleoplasmin to nuclear size-scaling. Related to Figure 4. **(A)** Stage 10 embryo extract and nuclei were mixed with egg extract cytosol at a 1:4 ratio (St10+Cyto) and supplemented with recombinant Npm2 (blue) or Npm-core (red) at the indicated concentrations or an equivalent volume of XB. After a 90-minute incubation, nuclei were fixed, spun onto coverslips, and stained with an antibody against the NPC (mAb414). Nuclear CS areas were measured for at least 650 nuclei per condition and normalized to the XB control. Data from two independent experiments are shown. Representative images are shown on the right. Error bars represent standard error. **(B)** Based on the nuclear growth curves shown in Figures 1C-E and 4A-B for ~0.1–0.4 nL droplets, the time to reach steady-state nuclear size was determined. The longer horizontal lines represent the means and the shorter horizontal lines correspond to the standard deviations. **(C)** An anti-Npm2 western blot was performed with 2 μl of egg extract cytosol (Cyto) and the indicated amounts of recombinant Npm2 protein. Band intensities were quantified and used to estimate the egg extract concentration of Npm2 as ~4.2 μM. **(D)** Based on reported values for average cell volumes at different developmental stages (Jevtic and Levy, 2015) and a measured Npm2 concentration of 4.2 μM that does not change significantly over early development (Fig. 4C), we estimated the per cell amount of Npm2 in different stages. We then plotted average nuclear volume (Jevtic and Levy, 2015) as a function of per cell Npm2 amounts and fit the data to a best-fit power regression curve. In our microinjection experiments, the total Npm2 concentration was increased from 4.2 μM to 7.7 μM, which corresponds to ~560 attomol of Npm2 per cell of a stage 10 embryo. Based on our regression curve, this amount of Npm2 would correspond to a nuclear volume of 3.5 pL if Npm2 was sufficient to account for nuclear-size scaling in the stage 8-10 developmental window. **(E)** One-cell *X. laevis* embryos were microinjected with equivalent volumes of XB or recombinant Npm2 protein (to increase the Npm2 concentration by 3.5 μM), and allowed to develop to stage 10-10.5. Nuclei isolated from embryos were fixed, spun onto coverslips, stained with an antibody against Npm2, and imaged with the same exposure time. 25 embryos on average were microinjected per condition. Representative images are shown. **(F)** One-cell *X. laevis* embryos were microinjected with equivalent volumes of XB, recombinant Npm2 protein (to increase the Npm2 concentration by 3.5 μM), importin-α-E mRNA (350 pg total), or recombinant Npm2 protein plus importin-α-E mRNA and allowed to develop to stage 10-10.5 without cycloheximide arrest. Importin-α-E is a phosphomimetic version of human importin α2 with reduced affinity for membranes (Levy and Heald, 2010; Wilbur and Heald, 2013). Nuclei isolated from embryos were fixed, spun onto coverslips, stained with an antibody against Npm2, and imaged with the same exposure time. Nuclear volumes and total nuclear Npm2 staining intensities were measured for at least 330 nuclei per condition per experiment. 25 embryos on average were microinjected per condition, and two independent experiments were performed. For the Npm2+importin α condition, representative images are shown and individual nuclear volume was plotted as a function of nuclear Npm2 staining intensity. For XB-microinjected embryos, nuclear Npm2 staining intensity was 2.05 ± 1.35 (average ± SD). In Figure 4F and 5C, the orange bars represent data for nuclei with Npm2 staining intensity values greater than one standard deviation above the XB control, therefore nuclei with Npm2 staining intensities greater than 2.05 + 1.35 = 3.4 (indicated by the open bracket on the x-axis in Figure S4F). ***, p≤0.001; ns, not significant. All error bars represent standard deviation, unless otherwise noted.

**Figure S5.**
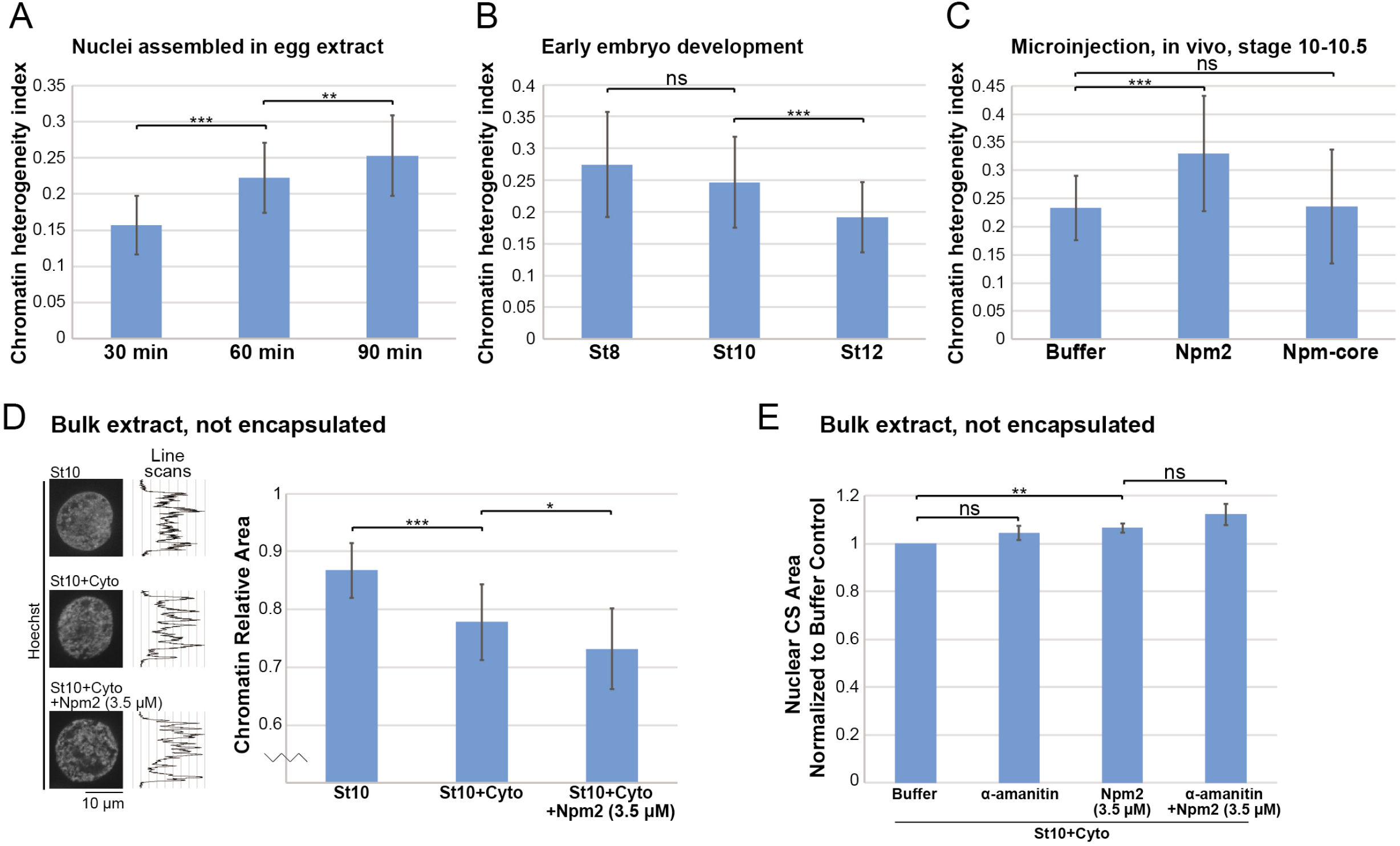
Chromatin topology measurements and demonstration that nucleoplasmin increases nuclear size independently of transcription. Related to Figure 5. **(A)** Nuclei were assembled de novo in interphasic *X. laevis* egg extract using demembranated *X. laevis* sperm as the chromatin source. After 30, 60, and 90 minutes of incubation, nuclei were fixed, spun down onto coverslips, and visualized with Hoechst. For each nucleus two Hoechst intensity line scans were acquired through the middle of the nucleus. The standard deviation of all intensity values along each line was determined and normalized to the average intensity to obtain a value we term the “chromatin heterogeneity index.” Larger values correspond to a more heterogeneous chromatin distribution. 54-68 line scans were quantified per condition (60 on average). **(B)** Nuclei from different stage embryo extracts were fixed, spun down onto coverslips, and visualized with Hoechst. For chromatin heterogeneity indexes, 22-79 line scans were quantified per condition (45 on average). **(C)** One-cell *X. laevis* embryos were microinjected with equivalent volumes of XB, recombinant Npm2 protein (to increase the Npm2 concentration by 3.5 μM), or recombinant Npm-core protein (3.5 μM final), allowed to develop to stage 10-10.5, and arrested in G2 with cycloheximide. Isolated nuclei were fixed, spun down onto coverslips, and visualized with Hoechst. For chromatin heterogeneity indexes, 34-52 line scans were quantified per condition (41 on average). **(D)** Stage 10 embryo extract and nuclei (St10) were mixed with egg extract cytosol at a 1:4 ratio (St10+Cyto) and supplemented with 3.5 μM recombinant Npm2 or an equivalent volume of XB. After a 1.5-hour incubation, nuclei were fixed, spun down onto coverslips, and visualized with Hoechst. Area occupied by Hoechst-staining chromatin was divided by total nuclear area to obtain chromatin relative area measurements. For each condition, 25-33 nuclei were quantified (28 nuclei on average). To the right of each representative Hoechst image is a Hoechst intensity line scan through the middle of the nucleus. **(E)** Stage 10 embryo extract and nuclei were mixed with egg extract cytosol at a 1:4 ratio (St10+Cyto) and supplemented with XB, 3.5 μM recombinant Npm2, and/or 100 μM α-amanitin, as indicated. After a 90-minute incubation, nuclei were fixed, spun down onto coverslips, and stained with an NPC antibody (mAb414). Nuclear CS areas were measured for at least 440 nuclei per condition and normalized to the XB control. Means and standard deviations from three independent experiments are shown. *, p≤0.05; **, p≤0.01; ***, p≤0.001; ns, not significant. All error bars represent standard deviation.

## MOVIE LEGENDS

**Movie 1. Nuclear growth in a 1 nL spherical droplet.** One stage 10 nucleus in stage 10 embryo extract was encapsulated in a ~1 nL spherical droplet and visualized by uptake of GFP-NLS. Images were acquired live at 30-second intervals with the same exposure time at room temperature. Imaging was performed for ~ 210 minutes. Related to Figure 1.

## Abbreviations

NE: nuclear envelope;
NPC: nuclear pore complex;
ER: endoplasmic reticulum;
NLS: nuclear localization signal;
PKC: protein kinase C;
CS: cross-sectional;
N/C: nuclear-to-cytoplasmic;
MBT: midblastula transition;
Npm2: nucleoplasmin

